# γδ T cells impair airway macrophage differentiation in lung adenocarcinoma

**DOI:** 10.1101/2023.09.14.557344

**Authors:** Ximena L. Raffo-Iraolagoitia, Amanda J. McFarlane, Bjorn Kruspig, Frédéric Fercoq, Judith Secklehner, Marco De Donatis, John B. G. Mackey, Robert Wiesheu, Sarah Laing, Ya-Ching Hsieh, Robin Shaw, Ryan Corbyn, Colin Nixon, Crispin Miller, Kristina Kirschner, Calum C. Bain, Daniel J. Murphy, Seth B. Coffelt, Leo M. Carlin

**Affiliations:** Cancer Research UK Beatson Institute, Glasgow, UK; School of Cancer Sciences, University of Glasgow, Glasgow, UK; Inflammation, Repair & Development, Imperial College London, London, UK; Institute for Regeneration and Repair, University of Edinburgh, Edinburgh, UK

## Abstract

Protecting mucosal barriers, γδ T cells hold promise for the development of new cancer immunotherapies. In mice, γδ T cells can largely be segregated into CD27^+^ and CD27^−^ cells, and their functions are modulated by interactions with surrounding cells. However, which cells communicate directly with γδ T cells in lung adenocarcinoma remains unknown. To address this, we combined flow cytometry, confocal microscopy, and scRNA-seq, using an autochthonous genetically engineered mouse model and different γδ T cell-deficient settings. We found that γδ T cells were increased in tumour-bearing lungs, with an altered phenotype. CD27^−^ and CD27^+^ γδ T cells differed in their localisation and interactions including their tropism for macrophages. Overall, we propose a model where CD27^+^ γδ T cells undermine the differentiation of tumour-associated macrophages into airway macrophages, fostering a negative outcome in lung adenocarcinoma. Determining its translatability to human health may offer new avenues for immunotherapeutic strategies.

**Summary:** Guardians of pulmonary homeostasis, γδ T cells remain enigmatic regarding their role in lung adenocarcinoma. Raffo-Iraolagoitia et al. report that a subset of γδ T cells impairs the differentiation of tumour-associated macrophages into airway macrophages, relevant for the outcome of lung adenocarcinoma.

**Graphical Abstract:** 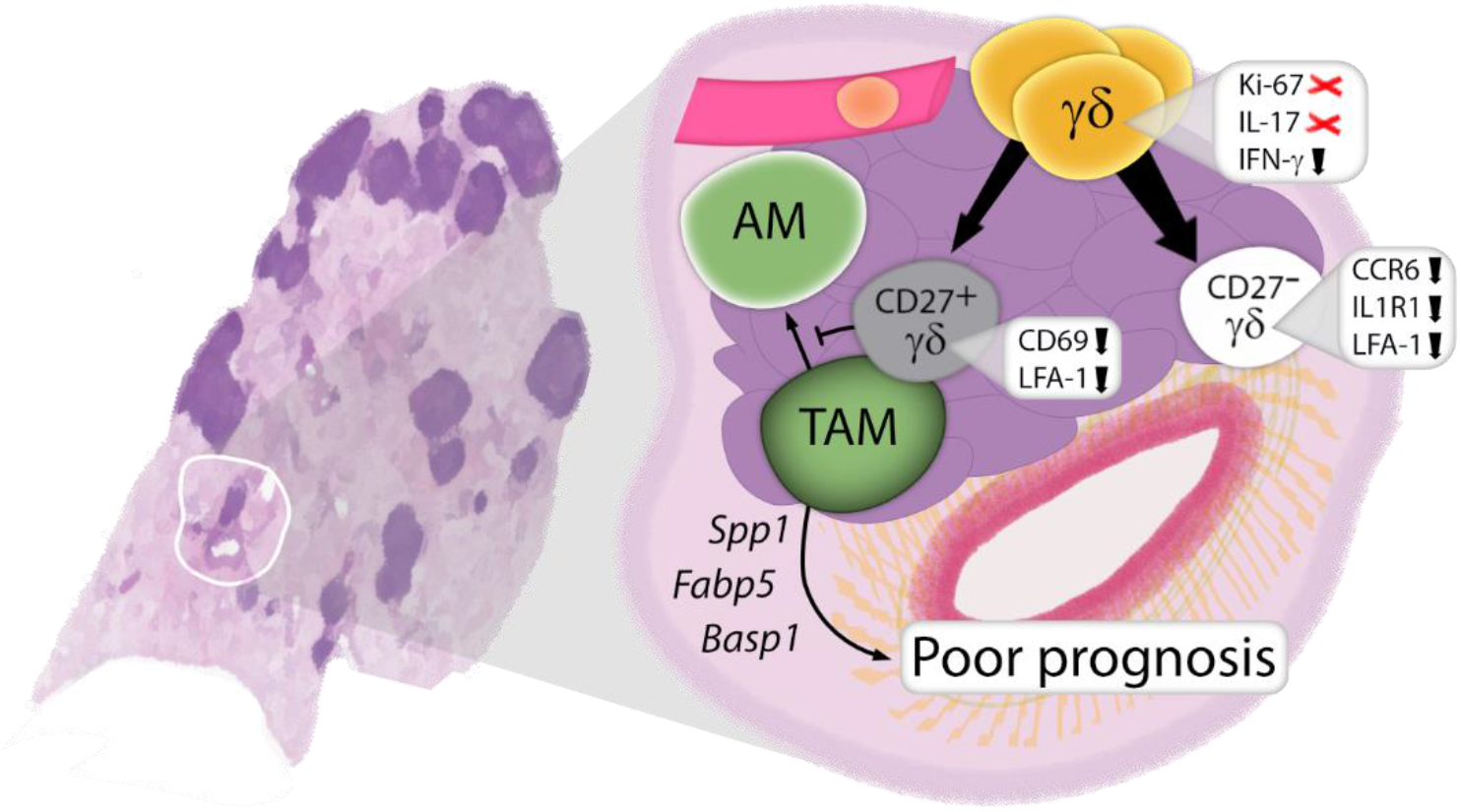

## INTRODUCTION

Pulmonary homeostasis relies on intricate cellular networks to secure gas exchange. In the lung, the immune response balances a high threshold upon innocuous particles and a robust clearance of invading pathogens through an interplay between pulmonary epithelial cells, tissue-resident immune cells, and leukocytes recruited from circulation (Lloyd and Marsland, 2017). Adding another layer of complexity, these cells are segregated into microanatomically distinct compartments such as parenchyma (including the respiratory bronchioles, alveolar ducts, and alveoli), perivascular adventitial cuffs (extracellular matrix-rich hubs for regional immune responses), and the intravascular leukocyte pool contained by the endothelium (Borger, 2020; Dahlgren and Molofsky, 2019). Comprehensively studying the changes that arise in these networks when epithelial cells become malignant might be the key to uncovering new targets for immunotherapies in lung cancer.

Gamma delta (γδ) T cells are unconventional T cells that provide protective and/or immunomodulatory functions at many barrier sites including the lung; and in non-small cell lung cancer (NSCLC) patients, the subset Vδ1 is associated with improved survival (Wu et al., 2022). In mice, γδ T cells consist of several subsets that express a variety of cytokines and other mediators of the immune response, and are generally subdivided into two populations depending on the expression of the cell surface marker CD27 (Ribot et al., 2020). In cancer, CD27^+^ γδ T cells are thought to play an anti-tumoural role due to their production of IFN-γ and cytotoxic ability (Silva-Santos et al., 2019), contrasting with their CD27^−^ counterparts which produce IL-17, supporting tumour development through a neutrophil-mediated mechanism (Coffelt et al., 2015; Edwards et al., 2023; Jin et al., 2019). However, our current understanding of their role in lung cancer is limited because their precise location within the pulmonary topology remains unknown.

Resting pulmonary γδ T cells have been shown to interact preferentially with myeloid cells, including macrophages (Wands et al., 2005). In the steady state lung, the macrophage compartment is heterogeneous comprising alveolar macrophages (AMs) and interstitial macrophages (IMs), which are transcriptionally and developmentally distinct. AMs are thought to maintain themselves autonomously in health through in situ self-renewal, however, upon inflammation, monocytes can also give rise to AMs (mo-AMs) (Misharin AV et al., 2017). In NSCLC, whereas AMs promote tumour formation at early times (Casanova-Acebes et al., 2021), these are replaced by mo-AMs which maintain an immunosuppressive role in anti-tumour immunity (Park et al., 2023). There have been several clinical trials attempting to target them in NSCLC such as Pexidartinib (NCT02452424), ARRY-382 (NCT02880371), Emactuzumab (NCT02323191), LY3022855 (Falchook et al., 2021; Dowlati et al., 2021), and AMG 80 (NCT02713529) (Razak et al., 2020; Papadopoulos et al., 2017), however no clinical benefit has been proven. In a mouse model of lung cancer, it was shown that AMs promote the expansion of γδ T cells (Jin et al., 2019). Whether, conversely, γδ T cells play a role in regulating AMs has not yet been explored.

In the context of lung adenocarcinoma (LuAd), we found that γδ T cells were increased in the lungs of tumour-bearing mice, where they displayed an altered phenotype. Spatial mapping revealed different tropisms for CD27^+^ and CD27^−^ γδ T cells within the LuAd microarchitecture. Our scRNA-seq data analysis indicated that γδ T cells, particularly in the CD27^+^ cluster, have increased probability of interaction with macrophages. We confirmed the occurrence of interactions between γδ T cells and tumour-associated macrophages (TAMs) by *ex-vivo* live and fixed tissue imaging. A pathogenic subset of tumour-associated AMs was reduced in CD27^+^ γδ T cell-deficient mice (transiently Vγ1-depleted) and *Tcrd* knockout (fully γδ T cell-deficient), but not in CD27^−^ γδ T cell-deficient KM mice (*TcrgV4/6* knockout). Overall, our findings reveal a potential pro-tumour role for the less abundant γδ T cell subset in the lung that has implications in the design of immunotherapies for LuAd as it directly influences the difficult to target tumour-promoting TAMs.

## RESULTS AND DISCUSSION

### Pulmonary γδ T cells increase in lung cancer

To characterise the response of γδ T cells to lung cancer, we used our well established autochthonous genetically engineered mouse model (GEMM). Here, a heterozygous point mutation from the endogenous *Kras* locus (*Kras*^*G12D*^) and modest overexpression of the oncogene MYC from the homozygous *Rosa26* locus were induced by intranasal delivery of an adenoviral vector containing Cre recombinase under the control of the surfactant protein C (*Spc*) promoter (herein referred to as SPC-Cre^+^ KM mice) (Kruspig et al., 2018). Eight weeks later, we analysed spleen, blood, and lungs by flow cytometry (**Fig. 1A**). At this time point, there were no clinical signs of disease, but early stage LuAds were evident in the lungs of SPC-Cre^+^ KM mice (**Fig. S1 A**). This was accompanied by an increase in lung leukocytes and reshaping of the immune landscape -with augmented monocytes, IMs, and AMs-specific to tumour-bearing mice, as SPC-Cre^+^ mice without the KM alleles showed no changes (**Fig. S1 B and C**). Although γδ T cells (**Fig. S1D**), remained a minority among other leukocytes, they were increased more than four-fold in SPC-Cre^+^ KM mice (**Fig. 1B**). However, we did not find evidence of increased in situ γδ T cell proliferation using Ki-67, or of their production of IL-17 or IFN-γ in tumour-bearing mice, in fact IFN-γ was reduced (**Fig. 1B**). Accordingly, we found a relative decrease in the CD27^+^ subpopulation over their CD27^−^ counterparts, compared with SPC-Cre^−^ KM mice (**Fig. 1B**). We next measured the expression of surface markers required for migration, activation, and adhesion in both subsets relative to their equivalents in SPC-Cre^−^ KM control mice (**Fig. 1C and S1E**). CCR6, whose downregulation is required for recruitment of γδ T cells to inflamed tissues (McKenzie et al., 2017), and IL1R1, critical for IL-17 production (Akitsu et al., 2015) were reduced in CD27^−^ but not in CD27^+^ γδ T cells. Conversely, the activation marker CD69 was reduced in CD27^+^ but not in CD27^−^ γδ T cells. Both subsets had reduced expression of LFA-1, an adhesion molecule involved in the formation of immune synapses that enhances cytotoxicity (Weng et al., 2021). Although small, none of these changes were mirrored in peripheral blood γδ T cells (**Fig. S1F)**, suggesting that the two subsets are regulated locally in the tumour microenvironment (TME). This is in contrast with previous findings in a GEMM driven by p53 where the role of γδ T cells recapitulated the classic circuit of IL-17 driven pro-tumour neutrophils (Jin et al., 2019). However, NSCLC patients have been reported to have a dysregulated γδ T cell repertoire (Bao et al., 2017), and never-smokers have more often unmutated TP53 (Clark et al., 2016). Taken together, this highlights the need to take into consideration a variety of mutational makeups when using preclinical models. As such, MYC is commonly dysregulated in cancer and has been shown to modulate the immune microenvironment (Casey et al., 2018; Kalkat et al., 2017; Kortlever et al., 2017; Mugarza et al., 2022).

**Fig. 1.**
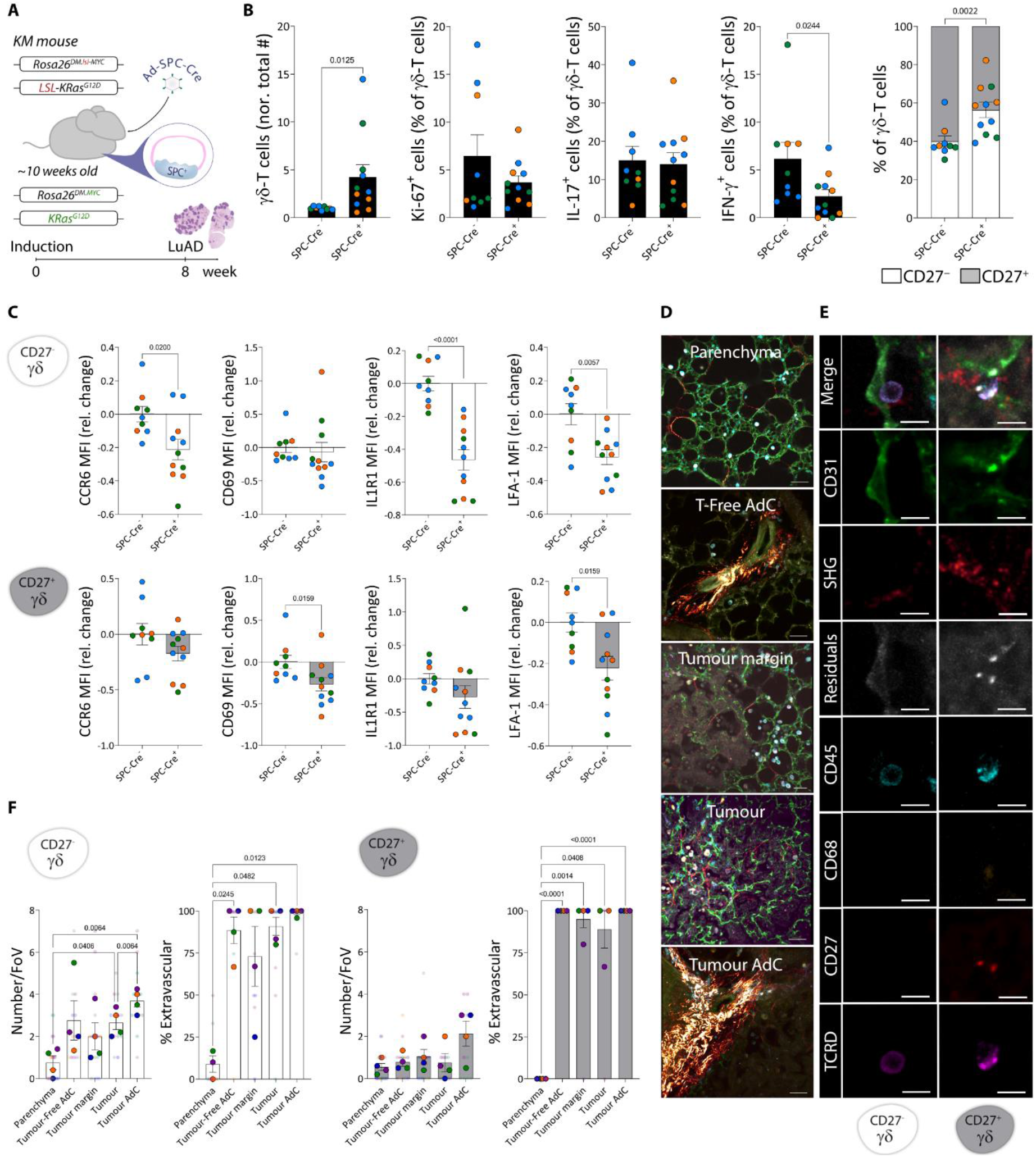
Pulmonary γδ T cells are increased, and change phenotype, in a GEMM of LuAd. **(A)** SPC-Cre^+^ KM mouse model of LuAd. Eight weeks after transgene activation, lungs were harvested for flow cytometry **(B-C)** or for confocal microscopy **(D-F). (B)** Left, total numbers of γδ T cells normalised to SPC-Cre^−^ KM controls. Right graphs, percentage of positive cells for Ki-67, IL-17, IFN-γ, and CD27. **(C)** Gated on CD27^−^ (white bars) and CD27^+^ (grey bars) γδ T cells, MFI relative change for CCR6, CD69, IL1R1, and LFA-1. Each dot represents a mouse, coloured by independent experiment. **(D)** Representative fields of view for each region type in precision-cut lung slices; 50μm scale bar. **(E)** Merged and individual channels for a representative CD27^−^ (left) and CD27^+^ (right) γδ T cell; 10μm scale bar. **(F)** Quantification of CD27^−^ and CD27^+^ γδ T cell mapping. Each colour represents a mouse, small dots for fields of view and big dots for means. Data are presented as mean ± SEM. Data were analysed by Mann-Whitney test **(B-C)**; one-way ANOVA or REML followed by Tukey’s multiple comparisons post-test **(F)**.

Next, we mapped the distribution of γδ T cells within the different topological compartments present in LuAd by confocal microscopy. We imaged five distinct regions based on the presence or absence of tumours and fibrillar collagen-rich adventitial cuffs (visualised by second harmonic generation), namely: parenchyma, tumour-free adventitial cuffs, tumour margin, tumour, and tumour-associated adventitial cuffs (**Fig. 1D**). In each region, we quantified γδ T cells as CD45^+^CD68^−^TCRδ^+^ and segregated these cells as either CD27^−^ or CD27^+^ (**Fig**.**1E**), determining their position relative to the vasculature (CD31). Whereas CD27^−^ γδ T cells were enriched in tumour regions, particularly surrounding adventitial cuffs, CD27^+^ γδ T cells did not show a striking tropism for a location. Both subsets were found in blood vessels in the parenchyma, however they mostly infiltrated the tissue in other regions (**Videos 1 and 2**). Thus, these results suggest that although γδ T cells display an altered phenotype, they are recruited to tumours.

### γδ T cells interact with TAMs

The interaction of γδ T cells with other cells in the lung microenvironment is poorly understood. Therefore, we sought to explore this network using CellChat (Jin et al., 2021). We performed total cell scRNA-seq of tumours microdissected under a stereomicroscope from SPC-Cre^+^ KM mouse lungs. Following the standard Seurat pipeline (Hao et al., 2021), we found seven clusters comprising epithelial cells, endothelial cells, and leukocytes (**Fig. 2A and S2A**). Of note, we could not identify a cluster of γδ T cells, even at higher resolutions, although this is not entirely surprising as γδ T cells in scRNA-seq experiments often do not cluster distinctly and appear embedded among more abundant lymphocytes such as CD8^+^ T cells and NK cells (Pizzolato et al., 2019). To predict interactions that naïve γδ T cells might have once recruited to the TME, we then leveraged these data with our scRNA-seq data from sorted pulmonary γδ T cells (Edwards et al., 2023). This dataset solely contained two clusters corresponding to CD27^−^ and CD27^+^ γδ T cells (**Fig. 2A and S2B**). Cellular communication modelling revealed all the interactions computed in the CellChat database that could be involved between the nine clusters together (**Fig. 2B**). Looking in detail at only those involving γδ T cells, we noticed that both CD27^−^ and CD27^+^ clusters were acting exclusively as “senders” in the significant pathways identified. The most probable “receivers” of outgoing signalling patterns from γδ T cells, particularly amongst those from the CD27^+^ subset, were macrophages through *Mrc1* (encoding CD206, a marker frequently associated with pro-tumour macrophages). Other putative receptor-ligand pairs included *Mif/Cd74* and *Ccl5/Ccr1* (**Fig. 2C**). In a chord diagram, using both subsets of γδ T cells as sources (bottom part of the ring), it was clear that most of the signalling outputs reaching macrophages (pink, lilac, and blue) were derived from CD27^+^ γδ T cells (orange arrows), and a minority of the *Ccl5* was supplied by CD27^−^ γδ T cells (violet arrow) (**Fig. 2 D**). Only a small fraction of the total interactions detected was received by clusters other than macrophages, possibly because γδ TCR-ligands are largely unknown. This *in silico* analysis allowed us to hypothesise that γδ T cells communicate with TAMs.

**Fig. 2.**
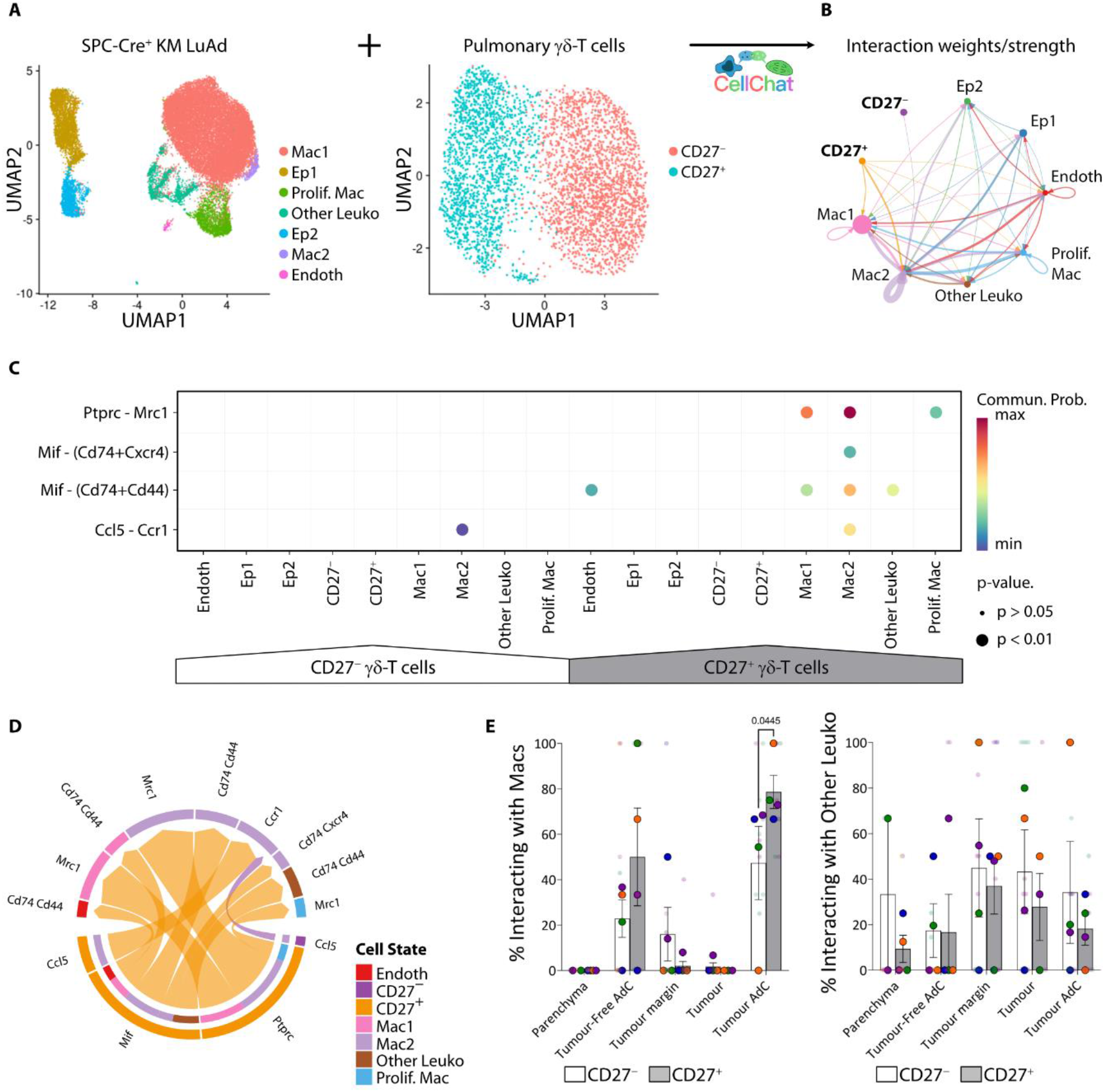
γδ T cells interact with TAMs. **(A)** ScRNA-seq data from microdissected SPC-Cre^+^ KM mouse tumours and naïve pulmonary γδ T cells (Edwards et al., 2023) were merged for CellChat analysis. **(B)** Aggregated cell-cell communication network visualised in a circle plot showing total interaction weights/strength between any two clusters. **(C)** Bubble plot showing all significant interactions from γδ T cells. **(D)** Chord diagram showing all significant interactions from γδ T cells. **(E)** Quantification by confocal microscopy of CD27^−^ (white) and CD27^+^ (grey) γδ T cells interacting with macrophages (CD45^+^CD68^+^, left panel) and other leukocytes (CD45^+^CD68^-^, right panel) grouped by different topologies in LuAd. Each colour represents a mouse, small dots for fields of view and big dots for means. Data are presented as mean ± SEM. Data were analysed by Mixed-effects Restricted Maximum Likelihood followed by uncorrected Fisher’s Least Significant Difference tests.

To address this hypothesis, we quantified the interactions of CD27^+^ γδ T cells and CD27^−^ γδ T cells with macrophages (CD45^+^CD68^+^) and other leukocytes (CD45^+^CD68^−^) within different topologies in LuAd (**Fig. 2E**). We used a semi-automated approach in Imaris manually verifying that cells were in contact and there were no other structures, such as endothelia, between them (**Video 3**). As predicted, a higher percentage of CD27^+^ than CD27^−^ γδ T cells were in close contact with macrophages, although only in tumour-associated adventitial cuffs. Whereas interactions with other leukocytes were also observed, these did not show any tropism or bias toward either subset of γδ T cells. As the quantification was performed in fixed tissue and could not discriminate whether the cells were dynamically interacting or just transient bystanders, we next explored live precision-cut lung slices (PCLS) to capture the behaviour of γδ T cells and TAMs in close contact. In this way, we could confirm the occurrence of long-lasting interactions of γδ T cells with TAMs in the tumour microenvironment (**Video 4**). Together, these data support an interplay between γδ T cells and TAMs in LuAd.

### TAMs are mainly composed of developmentally mature AMs and profibrotic AMs

TAMs are thought to promote tumour growth by a plethora of mechanisms, increasing the sources of heterogeneity of the different subsets of macrophages residing the lungs. Using scRNA-seq of SPC-Cre^+^ KM microdissected lung tumours, we found three clusters with transcriptional features of macrophages, including *Ear1, Car4*, and *Mrc1* (**Fig. 2a and S2C**). To address the heterogeneity of this population, we performed scRNA-seq of FACS isolated CD64^+^F4/80^+^ cells, including both AMs (SiglecF^+^CD11b^−^) and IMs (CD64^+^CD11b^+^), from control lungs (SPC-Cre^−^ KM) and from tumours microdissected under a stereomicroscope from SPC-Cre^+^ KM lungs (**Fig. S3A**). As most of the cells recovered were from SPC-Cre^+^ mice, we integrated our data with a reference dataset (McCowan et al., 2021) of pulmonary myeloid cells from control mice to enhance the numbers of cells from SPC-Cre^−^ lungs. Considering the annotation of the reference data set, we identified the predominant cluster in our data as AMs (**Fig. 3A**). Additionally, we found proliferating AMs and IMs, as well as dendritic cells and monocytes, but all these cells were scarce. We then built a heatmap for the AMs (cluster 0) with the 20 most differentially expressed genes across our conditions (SPC-Cre^−^ versus SPC-Cre^+^) (**Fig. 3B**). Interestingly, we noticed that 3 genes (*Basp1, Fabp5, Spp1*) out of the 20 markers were part of a consensus profibrotic signature driven by mo-AMs (Joshi et al., 2020). Indeed, tumour-associated AMs were enriched for this signature (**Fig. 3C and S3B**) which suggests they could be comprised of tissue-resident AMs and mo-AMs.

**Fig. 3.**
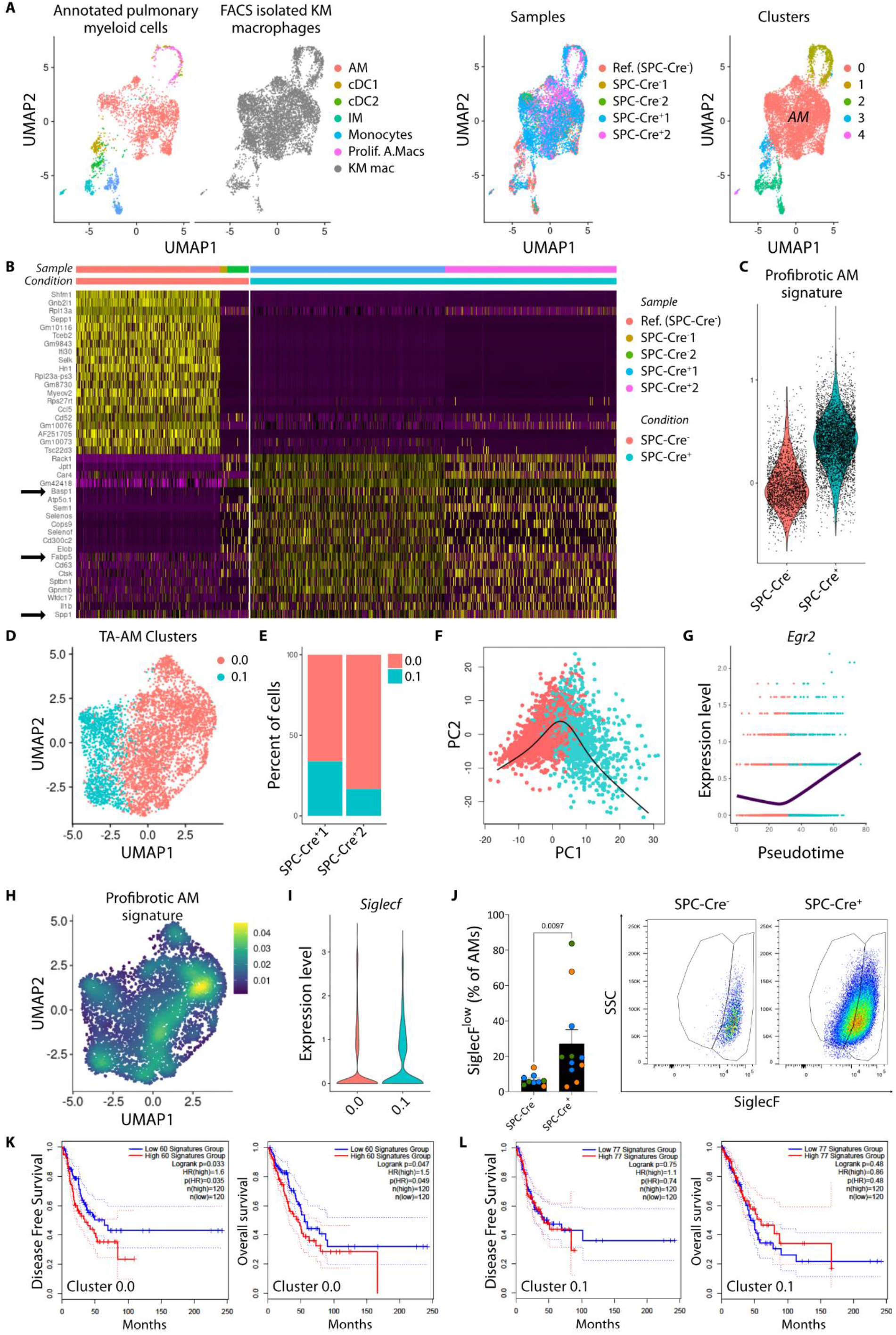
TAMs are mainly composed of developmentally mature AMs and profibrotic AMs. **(A)** UMAP plots of scRNA-seq from a reference pulmonary myeloid cell data set (McCowan et al., 2021) integrated with FACS isolated macrophages from microdissected KM lungs (SPC-Cre^−^ and SPC-Cre^+^) coloured by annotation (left), sample (centre) and cluster (right). **(B)** Heatmap showing the top 20 most differentially expressed genes in AMs (cluster 0) across conditions (SPC-Cre^−^ in red vs. SPC-Cre^+^ in blue). **(C)** Violin plot showing the expression of a consensus profibrotic AM signature (Joshi et al., 2020) by condition. AMs from SPC-Cre^+^ KM mice (TA-AM) were subsetted and re-clustered for further analysis **(D-I). (D)** UMAP plot of two subclusters of TA-AMs. **(E)** Stacked plot showing the percentage of each subcluster in two independent experiments. **(F)** Principal curve describing the global lineage structure with cluster 0.0 as starting point using Slingshot (Street et al., 2018) for pseudotime inference. **(G)** Expression level of *Egr2* over pseudotime from cluster 0.0 to cluster 0.1. **(H)** Density plot according to the profibrotic AM signature on UMAP dimensionality reduction for TA-AMs. **(I)** Violin plot showing the expression of Siglecf in TA-AMs clusters. **(J)** Percentage of SiglecF^low^ cells gated on AMs by flow cytometry. Each dot represents a mouse, coloured by experiment. Data are presented as mean ± SEM. Data were analysed by Mann-Whitney test. **(K-J)** Disease free and overall survival based on the expression of the signatures of Cluster 0.0 **(K)** or Cluster 0.1 **(L)** for LuAd patients in the GEPIA2 server (Tang et al., 2019).

Next, we re-clustered tumour-associated AMs revealing two transcriptionally distinct subpopulations with differences in 46 of the 50 GSEA Hallmark gene sets (**Fig. 3D, Fig. S3C**). These two clusters were similarly represented in both independent experiments (**Fig. 3E**). To explore whether there was a developmental difference between the clusters, we used Slingshot (Street et al., 2018) to fit a smooth trajectory using principal curves to describe the underlying lineage (**Fig. 3F**). When we analysed the expression of *Egr2*, a determinant of AM identity and function (McCowan et al., 2021), we observed that the expression of this transcription factor increased gradually over pseudotime from cluster 0.0 towards cluster 0.1 (**Fig. 3G**). Accordingly, the profibrotic mo-AM signature reported by Joshi, et al. (Joshi et al., 2020) was more prevalent in cluster 0.0 (**Fig. 3H**). Moreover, *Siglec5*, which encodes SiglecF and is considered the *de facto* marker of fully mature AMs, was expressed at lower levels by tumour-associated AM belonging to cluster 0.0 (**Fig. 3I**). Altogether, these findings indicate that tumour-associated AMs are comprised of two subsets with developmental differences. Indeed, we validated, by flow cytometry, the enrichment of a subpopulation of SiglecF^low^ AMs in SPC-Cre^+^ KM mice (**Fig. 3J**) previously described in models of acute lung injury where mo-AMs repopulate the lung after bleomycin treatment (Misharin AV et al., 2017) and influenza infection (Aegerter et al., 2020).

To assess the relevance to human health, we performed a survival analysis based on the signature of each cluster using the human orthologues of all differentially overexpressed genes in GEPIA2 (Tang et al., 2019). Whereas LuAd patients with high expression of the cluster 0.0 signature showed reduced disease free and overall survival (**Fig. 3K**), no significant stratification was observed when we used the cluster 0.1 signature (**Fig. 3L**). Overall, these results allow us to postulate that among tumour-associated AMs, there is a subset displaying a profibrotic signature that correlates with poor prognosis in LuAd. This finding complements previous research using intravenously injected cancer cells, where mo-AMs were associated with enhanced tumour spreading (Loyher et al., 2018).

### A subset of γδ T cells contribute to SiglecF^low^ AM accumulation

Given the physical interaction between CD27^+^ γδ T cells and TAMs, and the accumulation of SiglecF^low^ AMs in the lungs of tumour-bearing mice, we determined how macrophages, and specifically SiglecF^low^ AMs, are affected by the absence of γδ T cells. Thus, we crossed *Tcrd*^*—/—*^ mice with KM mice to generate cohorts of KM;*Tcrd*^*+/+*^ and KM;*Tcrd*^*—/—*^ mice. Eight weeks after allele activation, we analysed lungs as before (**Fig. 4A**). Whereas AMs were not affected in proportion or total number by the absence of γδ T cells, we observed that SPC-Cre^+^ KM; *Tcrd*^*—/—*^ mice had a decreased proportion of SiglecF^low^ AMs compared with SPC-Cre^+^ KM; *Tcrd*^*+/+*^ mice. Importantly, the change observed in the percentage of SiglecF^low^ AMs was independent of differences in tumour burden as these were equivalent at this time point (**Fig. 4B**).

**Fig. 4.**
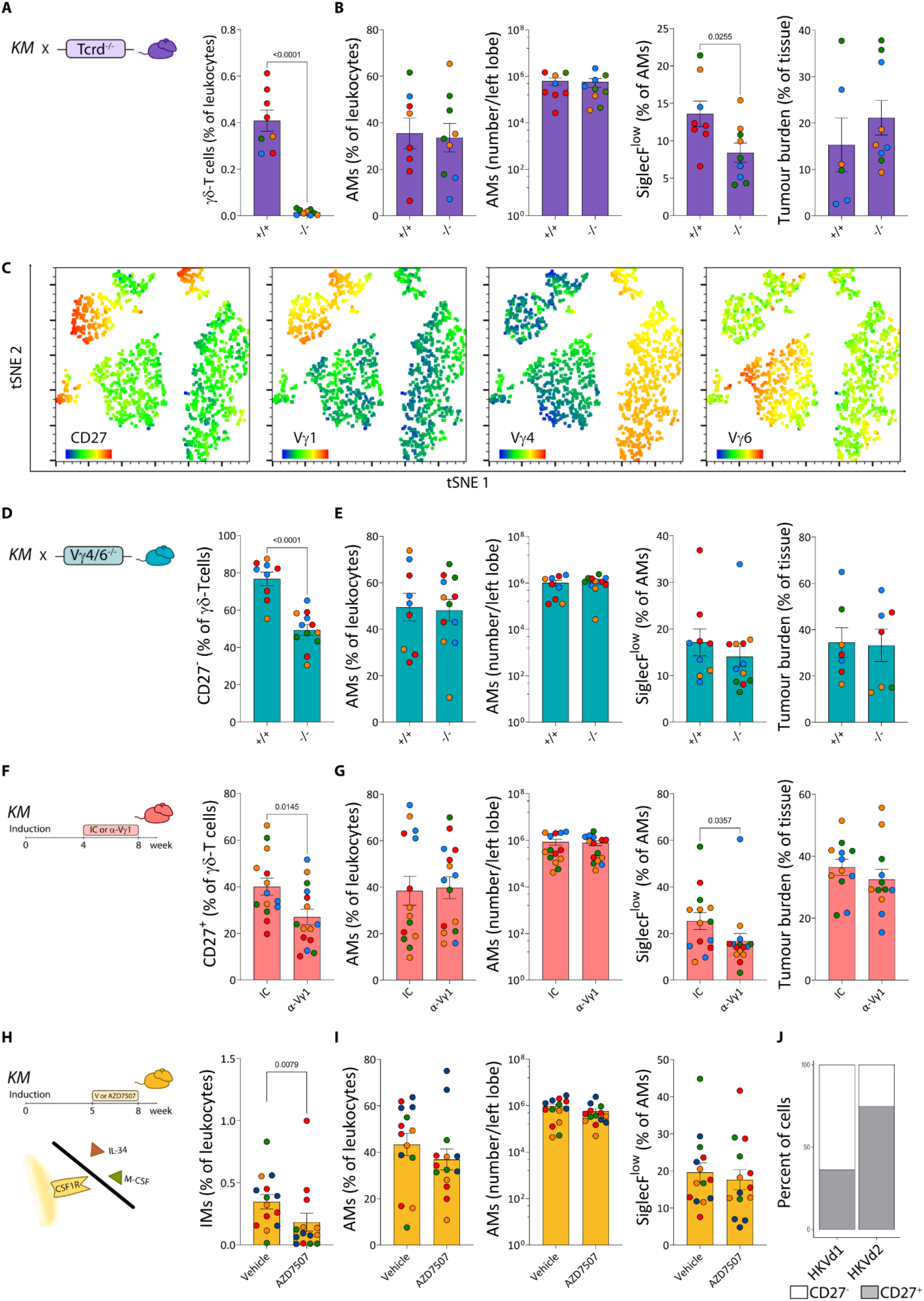
A subset of γδ T cells contribute to SiglecF^low^ AM accumulation. **(A)** KM mice were crossed with *Tcrd*^-/-^ mice and lungs were analysed 8 weeks post-induction. Percentage of γδ T cells in leukocytes (right). **(B)** Percentage of AMs in leukocytes, total number of AMs per left lobe, and percentage of SiglecF^low^ cells gated on AMs by flow cytometry. Tumour burden was assessed in whole lungs by histology. **(C)** tSNE dimensionality reduction using CD27, Vγ1, Vγ4 and Vγ6 as parameters on concatenated γδ T cells gated in SPC-Cre^+^ KM lung flow cytometry samples. **(D)** KM mice were crossed with *Vγ4/6*^-/-^ mice and lungs were analysed 8 weeks post-induction. Percentage of CD27^−^ cells gated on γδ T cells (right). **(E)** Percentage of AMs in leukocytes, total number of AMs per left lobe, and percentage of SiglecF^low^ cells gated on AMs by flow cytometry. Tumour burden was assessed in whole lungs by histology. **(F)** SPC-Cre^+^ KM mice were treated with isotype control (IC) or α-Vγ1 mAb for four weeks before analysis of lungs at week 8 post-induction. Percentage of CD27^+^ cells gated on γδ T cells (right). **(G)** Percentage of AMs in leukocytes, total number of AMs per left lobe, and percentage of SiglecF^low^ cells gated on AMs by flow cytometry. Tumour burden was assessed in whole lungs by histology. **(H)** SPC-Cre^+^ KM mice were treated with vehicle or AZD7507 for three weeks before analysis of lungs at week 8 post-induction. Percentage of IMs in leukocytes (right). **(I)** Percentage of AMs in leukocytes, total number of AMs per left lobe, and percentage of SiglecF^low^ cells gated on AMs by flow cytometry. Each dot represents a mouse, coloured by experiment. Data are presented as mean ± SEM. Data were analysed by t-test if Gaussian distribution **(A, B, D, and F)**, Mann-Whitney test otherwise **(G and H). (J)** Stacked plot showing the resemblance in scRNA-seq profiling between human γδ T cells split into δ1 and δ2 (Pizzolato et al., 2019), and mouse CD27^+^ and CD27^−^ γδ T cells (Edwards et al., 2023).

Beyond the more functional segregation of γδ T cells based on CD27 expression, γδ T cells can be further divided according to their γ-chain usage. Consistent with previous reports (Edwards et al., 2023; Ribot et al., 2020), in a t-SNE plot using CD27, Vγ1, Vγ4, and Vγ6 as parameters on concatenated γδ T cells from SPC-Cre^+^ KM mouse lungs, we found that most of the CD27^−^ γδ T cells are Vγ4^+^ or Vγ6^+^; and most of the CD27^+^ γδ T cells are Vγ1^+^ (**Fig. 4C**). To pinpoint whether there was a subset of γδ T cells responsible for the accumulation of SiglecF^low^ AMs, we targeted different Vγ chains either crossing KM mice with *Vγ4/6*^*—/—*^ mice or administering anti-Vγ1 antibody. In tumour-bearing mice induced as before, we observed an almost two-fold reduction in CD27^−^ γδ T cells, but AMs, SiglecF^low^ AMs, and tumour burden remained the same between KM;*Tcrd*^*+/+*^ and KM;*Vγ4/6*^*—/—*^ mice (**Fig. 4D and E**). Therefore, these data dismissed a role for Vγ4^+^ and Vγ6^+^ cells in the regulation of SiglecF^low^ AMs. Transient depletion of Vγ1 T cells in tumour-bearing KM mice reduced the proportion of CD27^+^ γδ T cells (**Fig. 4F**). While there were no changes in the proportion or total number of AMs between Vγ1^+^ cell-depleted and control mice, SPC-Cre^+^ KM mice receiving depleting antibodies displayed a decreased percentage of SiglecF^low^ AMs compared to control, with the same tumour burden (**Figure 4G**). These data indicate that Vγ1^+^ cells control AM differentiation in lung tumour-bearing mice.

As a CSF1-producing γδ T cell subset was recently described in a model of malaria (Mamedov et al., 2018), and tumours can employ similar mechanisms to parasitic diseases (Chulanetra and Chaicumpa, 2021; Rashidi et al., 2021; Babu Narasimhan et al., 2018) we next explored whether this molecule was involved in the accumulation of SiglecF^low^ AMs. Thus, we transiently blocked CSF1R in γδ T cell-sufficient mice and evaluated the AM phenotype as before. Consistent with their reliance on CSF1R, we found that administration of the CSF1R inhibitor AZD7507 led to a reduction in IMs (**Fig. 4H**). However, we found no changes amongst SiglecF-defined AMs (**Fig. 4I**). Taken together, these data suggest that the CD27^+^ γδ T cells that are largely Vγ1, but not the predominantly lung-resident CD27^−^-Vγ4/6 T cells, play a role in the increase of SiglecF^low^ AMs in a CSF1R-independent manner.

Finally, to further evaluate how these findings could potentially inform clinical research, we used publicly available scRNA-seq data from human γδ T cells (Pizzolato et al., 2019) to examine which human Vδ subsets better resemble the mouse CD27^−^ and CD27^+^ γδ T cells. To this end, we transformed the mouse γδ T cell scRNA-seq data set to human orthologues and queried their predicted identity in the human data set. We found that more than 50% of mouse CD27^−^ γδ T cells were labelled as human Vδ1, the subset that has been associated with increased survival in NSCLC (Wu et al., 2022), and the majority of the mouse CD27^+^ γδ T cells were more closely related to human Vδ2 (**Fig. 4J**).

In summary, we report that in our GEMM of LuAd, γδ T cells are recruited to tumours, where the CD27^+^ subset interacts preferentially with TAMs. We speculate that this interplay impairs AM differentiation as we demonstrated that Vγ1 cells mediate the accumulation of SiglecF^low^ AMs. These findings could be relevant to better understand paraneoplastic syndromes and common comorbidities such as pneumonia that often cost patients’ lives (Patel et al., 2020). Interestingly, whereas Vδ1 cells predict an extension in survival for NSCLC patients, there is not such benefit when considering all γδ T cells (Wu et al., 2022). This suggests that Vδ2 cells play a pro-tumour role that masks the protective role of Vδ1 cells. Determining whether human Vδ2 cells shape TAMs in a similar way as in our mouse model may offer new immunotherapeutic approaches in LuAd.

## MATERIALS AND METHODS

### Mouse models

The *LSL-KRas*^*G12D*^(Jackson et al., 2001), *Rosa26*^*DM*.*lsl-MYC*^ (Kruspig et al., 2018; RRID: IMSR_JAX:033805), *Tcrd*^*tm1Mom*^ (Itohara et al., 1993), and *Tcrg*^*tm1Iku*^ (Sunaga et al., 1997) alleles were previously described. All genetically modified mice were bred in house, maintained on a mixed FVB/N, C57BL/6 background, on a 12-hour light/dark cycle, and fed/watered ad libitum under specific pathogen-free conditions. Both males and females were included in approximately equal numbers and mice were randomly assigned to induction or treatment cohorts, balanced only for sex. KM mice were crossed with *Tcrd*^*—/—*^ mice (gifted to us from Adrian Hayday, The Francis Crick Institute, London, UK) or with *Vγ4/6*^*—/—*^ (gifted to us from Rebecca O’Brien, National Jewish Health, Denver, Colorado, USA). Control FVB/N mice were purchased from Charles River. All genotyping was performed by TransnetYX Inc.

Recombinant adenovirus Ad5mSPC-Cre, purchased from the University of Iowa gene therapy facility, was used to initiate lung adenocarcinoma by removing *lsl* cassettes from conditional alleles in alveolar type 2 *Spc* expressing cells. 1×10^8^ pfu Ad5mSPC-Cre viral particles were administered intranasally in 4 mM CaCl_2_ (Sigma) MEM medium (Thermo Fisher) to young adult (10-12 week-old) mice sedated with medetomidine and ketamine (Murphy et al., 2008). Atipamazole was administered to reverse anaesthesia.

For depletion of Vγ1^+^ cells, mice were injected intraperitoneally with anti-Vγ1 antibody (clone 2.11, BioXcell) beginning four weeks after allele activation with a single dose of 200 μg, followed by 100 μg twice per week thereafter. Control mice followed the same dosage regime with isotype control (polyclonal Armenian hamster IgG, BioXcell).

CSF1R inhibitor (AZD7507) was provided by Astra Zeneca under a research collaboration agreement with the CRUK Beatson Institute. Beginning five weeks after allele activation, AZD7507, dissolved in 0.5% HPMC and 0.1% Tween80 in distilled water, was administered at 100mg/kg by oral gavage twice daily for one week, and once per day thereafter. Control mice followed the same dosage regime with vehicle.

All mice were humanely killed by overdose of pentobarbital eight weeks after allele activation and tissues (spleen, blood and lungs) were harvested for downstream analysis. Procedures were performed in accordance with UK Home Office licence numbers 70/7950 and PE47BC0BF at the CRUK Beatson Institute.

### Tissue processing

Blood samples were withdrawn from the femoral artery using capillary action collection tubes with EDTA, and red blood cells were lysed using Ammonium-Chloride-Potassium buffer in 96-well V bottom plates. Spleens were mashed through a 70 μm filter with 5 mL DMEM with 5% FBS and 1% Penicillin/Streptomycin (Life Technologies). Before excising lungs, a small incision was made in the trachea, and a customised blunted needle was inserted. For some experiments, the left bronchus was tied with a suture and, subsequently, 0.6 mL of 2% low-melting point agarose in PBS was instilled slowly through the needle (1.1 mL in cases where whole lungs were inflated). Agarose inflated lungs were processed for confocal microscopy. Agarose-free left lobe or whole lungs were minced with scissors and incubated at 37°C with agitation in RPMI with 10% FBS, before mashing through a 70 μm filter. For scRNA-seq experiments, lungs were microdissected under a stereomicroscope. For the SPC-Cre^+^ KM tumour total cells dataset, enzymatic dissociation with 1 μg/ml Collagenase D (Roche) and 25 μg/mL DNAse I (ThermoFisher) was carried out using the gentleMACS Octo Dissociator (Miltenyi Biotec). Then, tumour suspensions were mashed through a 70 μm filter and live cells were enriched using debris removal solution (Miltenyi Biotec). For obtaining single cell suspensions for the KM macrophage datasets, microdissected tissue was processed in the same way than minced lungs. For histology, whole lungs were inflated through the trachea with 10% Neutral Buffered Formalin and fixed for 24h, before transferring to 70% Ethanol.

### Flow cytometry

For intracellular staining, single cell suspensions from lungs were incubated with Cell Stimulation Cocktail (eBioscience) in 96-well V bottom plates at 37°C for 3 hours. In one of the three experiments, a Percoll (GE Healthcare) gradient step was included before plating. After stimulation, cells were stained with a viability dye for 15 minutes at room temperature, followed by blocking with TruStain FCX PLUS (Biolegend), and then staining of surface markers in Brilliant stain buffer (BD Biosciences). After washing, cells were fixed and permeabilised in FOXP3 Transcription Factor Fixation/Permeabilisation solution (Thermo Fisher Scientific) following the manufacturer’s instructions. Antibodies for intracellular antigens were prepared in permeabilisation buffer and cells were incubated for 30 minutes at 4ºC. When assessing only surface markers, cells were similarly blocked, stained for viability, and then stained with a combination of antibodies detailed in **Table 1**. For scRNA-seq of macrophages, cells were sorted using a FACSAria II cell sorter (BD). For phenotyping, cells were fixed in 2% formaldehyde (VWR) and analysed using an LSR Fortessa analyser (BD) within a week. Just before acquisition, SPHERO ACCUCOUNT fluorescent beads (Spherotech) were added for quantification purposes. Data were analysed using FlowJo software (BD).

**Table 1:**
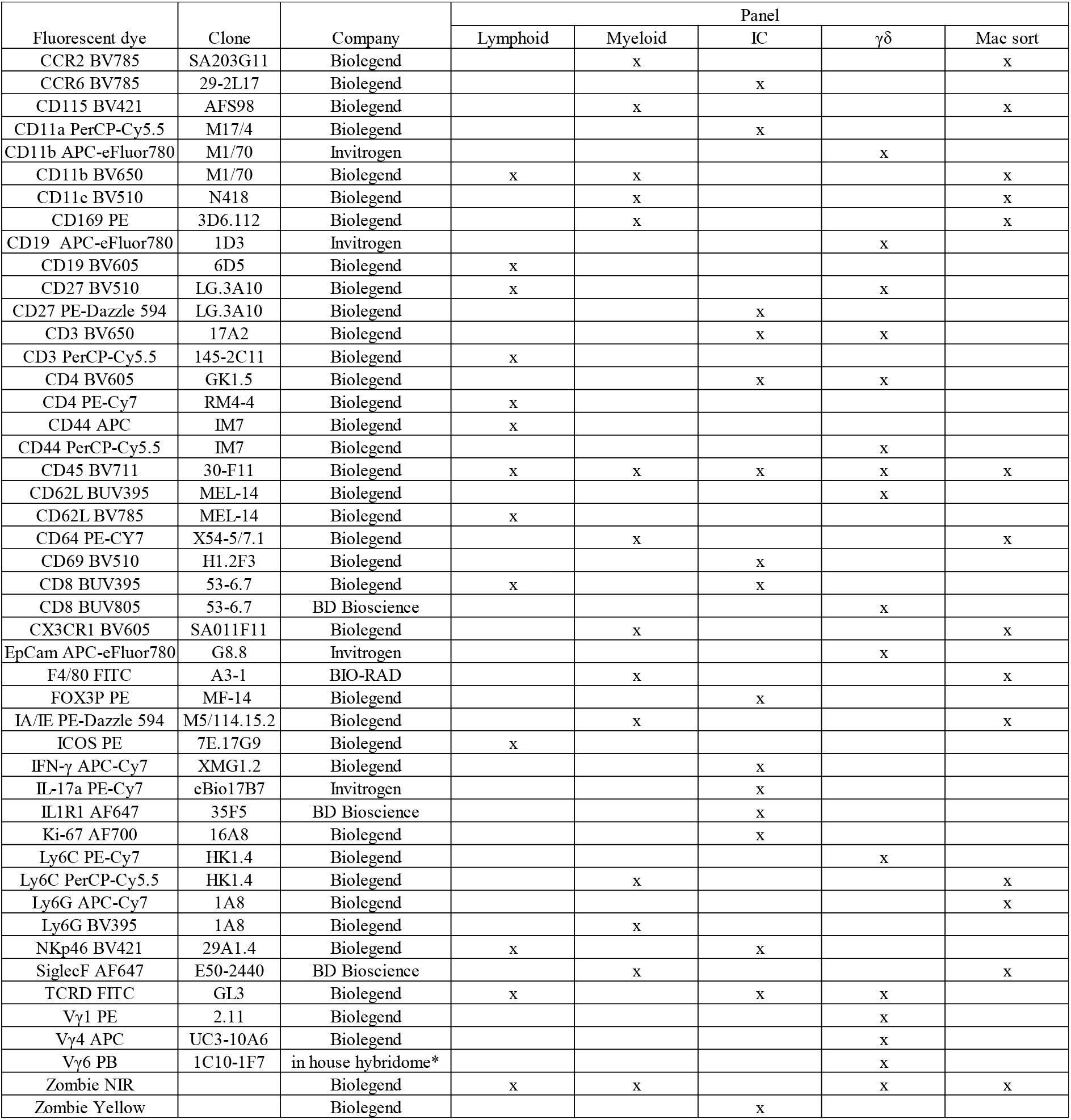
Flow cytometry reagents.

| Fluorescent dye | Clone | Company | Panel |  |  |  |  |
| --- | --- | --- | --- | --- | --- | --- | --- |
| | | | Lymphoid | Myeloid | IC | $\gamma\delta$ | Mac sort |
| CCR2 BV785 | SA203G11 | Biolegend |  | x |  |  | x |
| CCR6 BV785 | 29-2L17 | Biolegend |  |  | x |  |  |
| CD115 BV421 | AFS98 | Biolegend |  | x |  |  | x |
| CD11a PerCP-Cy5.5 | M17/4 | Biolegend |  |  | x |  |  |
| CD11b APC-eFluor780 | M1/70 | Invitrogen |  |  |  | x |  |
| CD11b BV650 | M1/70 | Biolegend | x | x |  |  | x |
| CD11c BV510 | N418 | Biolegend |  | x |  |  | x |
| CD169 PE | 3D6.112 | Biolegend |  | x |  |  | x |
| CD19 APC-eFluor780 | 1D3 | Invitrogen |  |  |  | x |  |
| CD19 BV605 | 6D5 | Biolegend | x |  |  |  |  |
| CD27 BV510 | LG.3A10 | Biolegend | x |  |  | x |  |
| CD27 PE-Dazzle 594 | LG.3A10 | Biolegend |  |  | x |  |  |
| CD3 BV650 | 17A2 | Biolegend |  |  | x | x |  |
| CD3 PerCP-Cy5.5 | 145-2C11 | Biolegend | x |  |  |  |  |
| CD4 BV605 | GK1.5 | Biolegend |  |  | x | x |  |
| CD4 PE-Cy7 | RM4-4 | Biolegend | x |  |  |  |  |
| CD44 APC | IM7 | Biolegend | x |  |  |  |  |
| CD44 PerCP-Cy5.5 | IM7 | Biolegend |  |  |  | x |  |
| CD45 BV711 | 30-F11 | Biolegend | x | x | x | x | x |
| CD62L BUV395 | MEL-14 | Biolegend |  |  |  | x |  |
| CD62L BV785 | MEL-14 | Biolegend | x |  |  |  |  |
| CD64 PE-Cy7 | X54-5/7.1 | Biolegend |  | x |  |  | x |
| CD69 BV510 | H1.2F3 | Biolegend |  |  | x |  |  |
| CD8 BUV395 | 53-6.7 | Biolegend | x |  | x |  |  |
| CD8 BUV805 | 53-6.7 | BD Bioscience |  |  |  | x |  |
| CX3CR1 BV605 | SA011F11 | Biolegend |  | x |  |  | x |
| EpCam APC-eFluor780 | G8.8 | Invitrogen |  |  |  | x |  |
| F4/80 FITC | A3-1 | BIO-RAD |  | x |  |  | x |
| FOX3P PE | MF-14 | Biolegend |  |  | x |  |  |
| IA/IE PE-Dazzle 594 | M5/114.15.2 | Biolegend |  | x |  |  | x |
| ICOS PE | 7E.17G9 | Biolegend | x |  |  |  |  |
| IFN-γ APC-Cy7 | XMG1.2 | Biolegend | | | x | | |
| IL-17a PE-Cy7 | eBio17B7 | Invitrogen |  |  | x |  |  |
| IL1R1 AF647 | 35F5 | BD Bioscience |  |  | x |  |  |
| Ki-67 AF700 | 16A8 | Biolegend |  |  | x |  |  |
| Ly6C PE-Cy7 | HK1.4 | Biolegend |  |  |  | x |  |
| Ly6C PerCP-Cy5.5 | HK1.4 | Biolegend |  | x |  |  | x |
| Ly6G APC-Cy7 | 1A8 | Biolegend |  |  |  |  | x |
| Ly6G BV395 | 1A8 | Biolegend |  | x |  |  |  |
| NKp46 BV421 | 29A1.4 | Biolegend | x |  | x |  |  |
| SiglecF AF647 | E50-2440 | BD Bioscience |  | x |  |  | x |
| TCRD FITC | GL3 | Biolegend | x |  | x | x |  |
| Vγ1 PE | 2.11 | Biolegend | | | | x | |
| Vγ4 APC | UC3-10A6 | Biolegend | | | | x | |
| Vγ6 PB | 1C10-1F7 | in house hybridome* | | | | x | |
| Zombie NIR |  | Biolegend | x | x |  | x | x |
| Zombie Yellow |  | Biolegend |  |  | x |  |  |

### Precision cut lung slices and confocal imaging

Fresh agarose-inflated lungs were sliced into 300μm thick sections on a vibrating microtome (Campden Ltd). For γδ T cell mapping, slices were blocked with TruStain FCX PLUS (Biolegend) and stained with anti TCRD AF647 (GL3, Biolegend), CD27 AF594 (LG.3A10, Biolegend), CD31 AF488 (390, Biolegend), and CD45 AF700 (30-F11, Biolegend) antibodies in complete medium (phenolred free DMEM supplemented with 1% FBS, Gibco) for 20 minutes at 37°C. After washing, slices were fixed in 4% formaldehyde in PBS (VWR) for 20 minutes at room temperature. Then, they were permeabilised and blocked in PBS with 0.3% Triton-X (Sigma), 10% normal goat serum (Sigma), 1% bovine serum albumin (Sigma), and 0.001% Sodium-Azide (VWR, UK); and subsequently stained with CD68 BV421 (FA-11, Biolegend) overnight at 4ºC. Finally, slices were mounted in Ce3D tissue clearing solution (Biolegend) and imaged on a an LSM 880 NLO upright confocal / multiphoton microscope (Carl Zeiss) equipped with a 32 channel Gallium arsenide phosphide (GaAsP) spectral detector using 20×/1 NA water immersion objective lens. Positions were selected based only on the topological features and excited simultaneously with 405, 488, 561 and 633 cw laser lines, and then with a pulsed two-photon laser tuned to 950 nm for second harmonic generation imaging of fibrillar collagen. For both acquisitions, signal was collected onto the 32 GaAsp detector linear array in lambda mode with a resolution of 8.9 nm over the visible spectrum. Spectral images were added together after linear unmixing with Zen software (Carl Zeiss) using reference spectra acquired from unstained tissues for autofluorescence and beads labelled with conjugated antibodies for each fluorophore. Alternatively, for live imaging, slices were stained with anti-CD31 AF647 (clone 390, Biolegend), TCRD AF488 (GL3, Biolegend) and CD11c AF594 (N418, Biolegend) antibodies, and Hoechst33342 (Thermo Scientific, UK) in complete medium for 20 minutes at 37°C. Slices were imaged on a LSM880 inverted confocal microscope with Airyscan detectors (Carl Zeiss) in a full incubation chamber at 37°C with 5% CO_2_. Airyscan time-lapse sequence processing was performed in Zen software (Carl Zeiss). Data were analysed with Imaris software (Bitplane, Oxford Instuments).

### scRNA-seq

Cells were resuspended in PBS containing 0.05% BSA and counted using a hemocytometer. A total of 40,000 cells were loaded onto each channel of Chromium Chip G using reagents from the 10x Chromium Single-Cell 3’ v3 Bead Kit and Library (10X Genomics) according to the manufacturer’s protocol. The libraries were analysed using the Bioanalyzer High Sensitivity DNA Kit (Aligent Technologies). scRNA-seq libraries were sequenced on the Illumina NovaSeq 6000 with paired-end 150-base reads. Quality checks and trimming on the raw scRNA-Seq fastq data files were done using FastQC (Andrews, 2022), FastP (Chen et al., 2018) and FastQ Screen (Wingett and Andrews, 2018). The creation of the transgenic reference genome and annotation was based on the GRCm38 version of the mouse genome and annotation (Cunningham et al., 2018). Alignment to the reference genome and aggregation was completed using 10x Genomics Cell Ranger version 3.1.0. Quality control, integration of data, clustering, marker gene identification and exploratory analysis was accomplished using the Seurat package version 4 (Hao et al., 2021) and the R environment version 4. Identification of high quality cells used the adaptive thresholds method as outlined in Orchestrating single-cell analysis with Bioconductor (Amezquita et al., 2020).

For the Seurat object of SPC-Cre^+^ KM tumours, 20 principal components were retained for clustering at 0.1 resolution. For the Seurat object of pulmonary mouse γδ T cell (ArrayExpress, #E-MTAB-10677), 15 principal components were retained for clustering at 0.2 resolution. Both datasets were appended with the Seurat command “merge”, then normalized and scaled (regressing out RNA counts) for subsequent analysis using CellChat (Jin et al., 2021).

The Seurat objects of KM macrophages and pulmonary myeloid cells (GSE181894) were integrated following normalization with SCTransform. Clustering was performed with 30 principal components at 0.1 resolution. Cluster 0 (AM) was subset and used to build a heatmap with the 20 top positive markers of “Condition”. The signature corresponding to “Profibrotic AM” was assessed using “AddModuleScore”. We re-clustered SPC-Cre^+^ AMs, taking 30 principal components at 0.15 resolution. For slingshot pipeline, cluster 0.0 was set as start. Differentially upregulated genes were obtained using “FindAllMarkers” with the test MAST. These genes were converted to human orthologous symbols using the library nichenetr (Browaeys et al., 2020) to interrogate GEPIA2 database (Tang et al., 2019). The R package Escape version 3.17 was used to calculate ssGSEA (Barbie et al., 2009) using gene sets from the Molecular Signatures Database (Subramanian et al., 2005). The Mann Whitney U test was used to determine significant differences. The ReactomeGSA package (Griss et al., 2020) was used to perform GSVA (Hänzelmann et al., 2013) using data from the Reactome pathway knowledgebase (Gillespie et al., 2021).

We used nichenetr to convert the pulmonary mouse γδ T cell dataset into human orthologous, and then we created a Seurat object filtering the expression matrix for genes which a human counterpart was available. Clustering was performed with 15 principal components at 0.2 resolution. We used this Seurat object as a query to assess the resemblance of CD27^−^ and CD27^+^ γδ T cells with human γδ T cell subsets from the GSE128223 data set.

All computational analysis was documented at each stage using MultiQC (Ewels et al., 2016), Jupyter Notebooks (Kluyver et al., 2016) and/or R Notebooks.

### Histology

Lungs were loaded into labelled histology processing cassettes and processed through a series of graded ethanol (70%, 90%, 95, 100% x3), 3 changes of xylene and 3 changes of histology wax applied under pressure. This was performed using an overnight tissue processing cycle on an Epredia excelsior tissue processor. Following this, the tissue was orientated and embedded in histology wax to create a formalin fixed paraffin embedded (FFPE) block. 4μm sections were cut from each FFPE block, placed onto a glass slide, and the sections ovened at 60°C for 2 hours.

Haematoxylin and eosin (H&E) staining was performed on a Leica autostainer (ST5020). Sections were dewaxed in xylene, taken through graded ethanol solutions and stained with Haem Z (RBA-4201-00A, CellPath) for 13 mins. Sections were washed in water, differentiated in 1% acid alcohol, washed, and nuclei blued in Scott’s tap water substitute (in-house). After washing with tap water sections were placed in Putt’s Eosin (in-house) for 3 minutes. The H&E sections were washed in tap water, dehydrated through graded ethanol solutions, and placed in xylene. The stained sections were coverslipped from xylene using DPX mountant (SEA-1300-00A, CellPath). Once the DPX mountant had solidified, the glass slides were loaded onto a Leica Aperio AT2 slide scanner, and digital images captured at x20 magnification. Tumour burden was determined for each mouse using QuPath (Bankhead et al., 2017) as the % of lung tissue occupied by tumours, averaging H&E slides from three paraffin-embedded sections at 400 μm intervals.

### Statistical analysis

Normalised total number of cells was calculated as the total number of cells in a sample, divided by the average total number of cells in control samples for each independent experiment; this allowed us to combine two experiments where whole lungs were processed for flow cytometry with one experiment where the right lobes were used for imaging. MFI relative change was calculated as the difference between MFI of the gated population and the average MFI of the same gating in control samples, divided by the average MFI of the same gating in control samples, for each independent experiment. Statistical analysis was performed using GraphPad Prism (version 9.5.1) as detailed in figure legends.

## Supplemental material

The supplementary material shows the immunephenotype of KM mice (**Fig. S1**), density and violin plots for the genes used to annotate scRNA-seq clusters (**Fig. S2**), and data pertinent to the scRNA-seq experiment on KM macrophages (**Fig. S3**). **Video 1** shows a 3D view of the parenchyma zooming-in on intravascular γδ T cells. **Video 2** shows a 3D view of a lung tumour zooming-in on an infiltrating γδ T cell. **Video 3** shows a 3D view of the lung tumour-associated adventitial cuff zooming-in on an interacting γδ T cell. **Video 4** shows long-lasting interactions between γδ T cells and AMs.

## Acknowledgments

We would like to thank Catherine Winchester and Kirsteen Campbell for critical reading of the manuscript. We would like to thank the core services and advanced technologies at the CRUK Beatson Institute with particular thanks to the Biological Services Unit, Bioinformatics and the Beatson Advanced Imaging Resource. We would also like to thank, Ed W Roberts, Sarah C. Edwards, Anna Kilbey, Olympia Hardy, Ross Laidlaw, Alexandrina Pancheva, and all past and present members of the Leukocyte Dynamics Group for helpful discussions during the project.

## Author contributions

Conceptualization, project administration, and writing-original draft by XLRI and LMC. Investigation by XLRI, AJM, BK, FF, JS, MDD, JBGM, RW, SL, YCH, and CN. Visualization by XLRI. Formal analysis by XLRI, RS, and RC. Software by XLRI and RS. Data curation by RS. Resources by CCB, KK, DJM, SBC, and LMC. XLRI, CN, CM, KK, CCB, DJM, SBC, and LMC: supervision. Writing-review & editing by all authors.

## Data and materials availability

The accession number of the scRNA-seq data sets will be available after peer-review of the manuscript. All other data are available at reasonable request by contacting the corresponding authors.

## References and Notes

Aegerter, H., J. Kulikauskaite, S. Crotta, H. Patel, G. Kelly, E.M. Hessel, M. Mack, S. Beinke, and A. Wack. 2020. Influenza-induced monocyte-derived alveolar macrophages confer prolonged antibacterial protection. Nat Immunol. 21:145–157. doi:10.1038/s41590-019-0568-x.

Akitsu, A., H. Ishigame, S. Kakuta, S.-H. Chung, S. Ikeda, K. Shimizu, S. Kubo, Y. Liu, M. Umemura, G. Matsuzaki, Y. Yoshikai, S. Saijo, and Y. Iwakura. 2015. IL-1 receptor antagonist-deficient mice develop autoimmune arthritis due to intrinsic activation of IL-17-producing CCR2+ Vgamma6+ gamma delta T cells. doi:10.1038/ncomms8464.

Amezquita, R.A., A.T.L. Lun, E. Becht, V.J. Carey, L.N. Carpp, L. Geistlinger, F. Marini, K. Rue-Albrecht, D. Risso, C. Soneson, L. Waldron, H. Pagès, M.L. Smith, W. Huber, M. Morgan, R. Gottardo, and S.C. Hicks. 2020. Data infrastructure Orchestrating single-cell analysis with Bioconductor. Nat Methods. 17:137–145. doi:10.1038/s41592-019-0654-x.

Andrews, S. 2022. FastQC A Quality Control tool for High Throughput Sequence Data. Babraham Bioinformatics.

Babu Narasimhan, P., L. Akabas, S. Tariq, N. Huda, S. Bennuru, H. Sabzevari, R. Hofmeister, T.B. Nutman, and R. Tolouei Semnani. 2018. Similarities and differences between helminth parasites and cancer cell lines in shaping human monocytes: Insights into parallel mechanisms of immune evasion. PLoS Negl Trop Dis. 12. doi:10.1371/journal.pntd.0006404.

Bankhead, P., M.B. Loughrey, J.A. Fernández, Y. Dombrowski, D.G. Mcart, P.D. Dunne, S. Mcquaid, R.T. Gray, L.J. Murray, H.G. Coleman, J.A. James, M. Salto-Tellez, and P.W. Hamilton. 2017. QuPath: Open source software for digital pathology image analysis. Sci Rep. 7. doi:10.1038/s41598-017-17204-5.

Bao, Y., L. Guo, and J. Mo. 2017. Characterization of γδ T cells in patients with non-small cell lung cancer. Oncol Lett. 14:1133. doi:10.3892/OL.2017.6191.

Barbie, D.A., P. Tamayo, J.S. Boehm, S.Y. Kim, S.E. Moody, I.F. Dunn, A.C. Schinzel, P. Sandy, E. Meylan, C. Scholl, S. Fröhling, E.M. Chan, M.L. Sos, K. Michel, C. Mermel, S.J. Silver, B.A. Weir, J.H. Reiling, Q. Sheng, P.B. Gupta, R.C. Wadlow, H. Le, S. Hoersch, B.S. Wittner, S. Ramaswamy, D.M. Livingston, D.M. Sabatini, M. Meyerson, R.K. Thomas, E.S. Lander, J.P. Mesirov, D.E. Root, D.G. Gilliland, T. Jacks, and W.C. Hahn. 2009. Systematic RNA interference reveals that oncogenic KRAS-driven cancers require TBK1. Nature. doi:10.1038/nature08460.

Borger, J.G. 2020. Spatiotemporal Cellular Networks Maintain Immune Homeostasis in the Lung. EMJ Respir. 8:108–119.

Browaeys, R., W. Saelens, and Y. Saeys. 2020. NicheNet: modeling intercellular communication by linking ligands to target genes. Nat Methods. 17:159–162. doi:10.1038/s41592-019-0667-5.

Casanova-Acebes, M., E. Dalla, A.M. Leader, J. LeBerichel, J. Nikolic, B.M. Morales, M. Brown, C. Chang, L. Troncoso, S.T. Chen, A. Sastre-Perona, M.D. Park, A. Tabachnikova, M. Dhainaut, P. Hamon, B. Maier, C.M. Sawai, E. Agulló-Pascual, M. Schober, B.D. Brown, B. Reizis, T. Marron, E. Kenigsberg, C. Moussion, P. Benaroch, J.A. Aguirre-Ghiso, and M. Merad. 2021. Tissue-resident macrophages provide a pro-tumorigenic niche to early NSCLC cells. Nature. 595. doi:10.1038/s41586-021-03651-8.

Casey, S.C., V. Baylot, and D.W. Felsher. 2018. The MYC oncogene is a global regulator of the immune response. Blood. 131:2007–2015.

Chen, S., Y. Zhou, Y. Chen, and J. Gu. 2018. fastp: an ultra-fast all-in-one FASTQ preprocessor. Bioinforma Oxf Engl. 34:i884–90. doi:10.1093/bioinformatics/bty560.

Chulanetra, M., and W. Chaicumpa. 2021. Revisiting the Mechanisms of Immune Evasion Employed by Human Parasites. Front Cell Infect Microbiol. 11. doi:10.3389/fcimb.2021.702125.

Clark, G., H. Donninger, S. Piscuoglio, A.R. Halvorsen, L. Silwal-Pandit, L.A. Meza-Zepeda, D. Vodak, P. Vu, C. Sagerup, E. Hovig, O. Myklebost, A.-L. Børresen-Dale, O.T. Brustugun, and Å. Helland. 2016. TP53 Mutation Spectrum in Smokers and Never Smoking Lung Cancer Patients. Front Genet. 7. doi:10.3389/fgene.2016.00085.

Coffelt, S.B., K. Kersten, C.W. Doornebal, J. Weiden, K. Vrijland, C.-S. Hau, N.J. M Verstegen, M. Ciampricotti, L.J. A C Hawinkels, J. Jonkers, and K.E. de Visser. 2015. IL-17-producing cd T cells and neutrophils conspire to promote breast cancer metastasis. Nature. 522:345–348. doi:10.1038/nature14282.

Cunningham, F., P. Achuthan, W. Akanni, J. Allen, M.R. Amode, I.M. Armean, R. Bennett, J. Bhai, K. Billis, S. Boddu, C. Cummins, C. Davidson, J. Dodiya, A. Gall, C. García Girón, G. Girón, L. Gil, T. Grego, L. Haggerty, E. Haskell, T. Hourlier, O.G. Izuogu, S.H. Janacek, T. Juettemann, M. Kay, M.R. Laird, I. Lavidas, Z. Liu, J.E. Loveland, J. Joś, J.C. Marugán, M. Marugán, T. Maurel, A.C. Mcmahon, B. Moore, J. Morales, J.M. Mudge, M. Nuhn, D. Ogeh, A. Parker, A. Parton, M. Patricio, A. Imran, A. Salam, B.M. Schmitt, H. Schuilenburg, D. Sheppard, H. Sparrow, E. Stapleton, M. Szuba, K. Taylor, G. Threadgold, A. Thormann, A. Vullo, B. Walts, A. Winterbottom, A. Zadissa, M. Chakiachvili, A. Frankish, S.E. Hunt, M. Kostadima, N. Langridge, F.J. Martin, M. Muffato, E. Perry, M. Ruffier, D.M. Staines, S.J. Trevanion, B.L. Aken, A.D. Yates, D.R. Zerbino, and P. Flicek. 2018. Ensembl 2019. Nucleic Acids Res. 47:745–751. doi:10.1093/nar/gky1113.

Dahlgren, M.W., and A.B. Molofsky. 2019. Adventitial Cuffs: Regional Hubs for Tissue Immunity. Trends Immunol. 40:877–887. doi:10.1016/j.it.2019.08.002.

Dowlati, A., R. Donald Harvey, R.D. Carvajal, O. Hamid, S.J. Klempner, J. Sae Wook Kauh, D.A. Peterson, D. Yu, S.C. Chapman, A.M. Szpurka, M. Carlsen, T. Quinlan, and R. Wesolowski. 2021. LY3022855, an anti–colony stimulating factor-1 receptor (CSF-1R) monoclonal antibody, in patients with advanced solid tumors refractory to standard therapy: phase 1 dose-escalation trial. Invest New Drugs. 39:1057–1071. doi:10.1007/s10637-021-01084-8/Published.

Edwards, S.C., A. Hedley, W.H. Hoevenaar, R. Wiesheu, T. Glauner, A. Kilbey, R. Shaw, K. Boufea, N. Batada, S. Hatano, Y. Yoshikai, K. Blyth, C. Miller, K. Kirschner, and S.B. Coffelt. 2023. PD-1 and TIM-3 differentially regulate subsets of mouse IL-17A-producing γδ T cells. J Exp Med. 220. doi:10.1084/jem.20211431.

Ewels, P., M. Ns Magnusson, S. Lundin, and M.K. Aller. 2016. MultiQC: summarize analysis results for multiple tools and samples in a single report. Bioinformatics. 32. doi:10.1093/bioinformatics/btw354.

Falchook, G.S., M. Peeters, S. Rottey, L.Y. Dirix, R. Obermannova, J.E. Cohen, R. Perets, R. Shapira Frommer, T.M. Bauer, J.S. Wang, R.D. Carvajal, J. Sabari, S. Chapman, W. Zhang, B. Calderon, D.A. Peterson, and G.S. Falchook GeraldFalchook. 2021. PHASE I STUDIES A phase 1a/1b trial of CSF-1R inhibitor LY3022855 in combination with durvalumab or tremelimumab in patients with advanced solid tumors. Invest New Drugs. 39:1284–1297. doi:10.1007/s10637-021-01088-4/Published.

Gillespie, M., B. Jassal, R. Stephan, M. Milacic, K. Rothfels, A. Senff-Ribeiro, J. Griss, C. Sevilla, L. Matthews, C. Gong, C. Deng, T. Varusai, E. Ragueneau, Y. Haider, B. May, V. Shamovsky, J. Weiser, T. Brunson, N. Sanati, L. Beckman, X. Shao, A. Fabregat, K. Sidiropoulos, J. Murillo, G. Viteri, J. Cook, S. Shorser, G. Bader, E. Demir, C. Sander, R. Haw, G. Wu, L. Stein, H. Hermjakob, and P. D’eustachio. 2021. The reactome pathway knowledgebase 2022. Nucleic Acids Res. 50. doi:10.1093/nar/gkab1028.

Griss, J., G. Viteri, K. Sidiropoulos, V. Nguyen, A. Fabregat, and H.H. Correspondence. 2020. ReactomeGSA-Efficient Multi-Omics Comparative Pathway Analysis. Mol Cell Proteomics. 19:2115–2124. doi:10.1074/mcp.TIR120.002155.

Hänzelmann, S., R. Castelo, and J. Guinney. 2013. GSVA: gene set variation analysis for microarray and RNA-seq data. BMC Bioinformatics. 14:7. doi:10.1186/1471-2105-14-7.

Hao, Y., S. Hao, E. Andersen-Nissen, W.M. Mauck, S. Zheng, A. Butler, M.J. Lee, A.J. Wilk, C. Darby, M. Zager, P. Hoffman, M. Stoeckius, E. Papalexi, E.P. Mimitou, J. Jain, A. Srivastava, T. Stuart, L.M. Fleming, B. Yeung, A.J. Rogers, J.M. McElrath, C.A. Blish, R. Gottardo, P. Smibert, and R. Satija. 2021. Integrated analysis of multimodal single-cell data. Cell. 184:3573–3587.e29. doi:10.1016/J.CELL.2021.04.048.

Itohara, S., P. Mombaerts, J. Lafaille, J. Iacomini, A. Nelson, A.R. Clarke, M.L. Hooper, A. Farr, and S. Tonegawa. 1993. T cell receptor δ gene mutant mice: Independent generation of αβ T cells and programmed rearrangements of γδ TCR genes. Cell. 72:337–348. doi:10.1016/0092-8674(93)90112-4.

Jackson, E.L., N. Willis, K. Mercer, R.T. Bronson, D. Crowley, R. Montoya, T. Jacks, and D.A. Tuveson. 2001. Analysis of lung tumor initiation and progression using conditional expression of oncogenic K-ras. Genes Dev. 15:3243–8. doi:10.1101/gad.943001.

Jin, C., G.K. Lagoudas, C. Zhao, P.C. Blainey, J.G. Fox, and T. Jacks. 2019. Commensal Microbiota Promote Lung Cancer Development via gamma delta T Cells. Cell. 176:998–1013. doi:10.1016/j.cell.2018.12.040.

Jin, S., C.F. Guerrero-Juarez, L. Zhang, I. Chang, R. Ramos, C.-H. Kuan, P. Myung, M. V Plikus, and Q. Nie. 2021. Inference and analysis of cell-cell communication using CellChat. Nat Commun. 12. doi:10.1038/s41467-021-21246-9.

Joshi, N., S. Watanabe, R. Verma, R.P. Jablonski, C.-I. Chen, P. Cheresh, N.S. Markov, P.A. Reyfman, A.C. Mcquattie-Pimentel, L. Sichizya, Z. Lu, R. Piseaux-Aillon, D. Kirchenbuechler, A.S. Flozak, C.J. Gottardi, C.M. Cuda, H. Perlman, M. Jain, D.W. Kamp, G.R. Scott Budinger, and A. V Misharin. 2020. A spatially restricted fibrotic niche in pulmonary fibrosis is sustained by M-CSF/M-CSFR signalling in monocyte-derived alveolar macrophages. Eur Respir J. 55. doi:10.1183/13993003.00646-2019.

Kalkat, M., J. De Melo, K.A. Hickman, C. Lourenco, C. Redel, D. Resetca, A. Tamachi, W.B. Tu, and L.Z. Penn. 2017. MYC Deregulation in Primary Human Cancers. Genes (Basel). 8:151. doi:10.3390/genes8060151.

Kluyver, T., B. Ragan-Kelley, F. Pérez, B. Granger, M. Bussonnier, J. Frederic, K. Kelley, J. Hamrick, J. Grout, S. Corlay, P. Ivanov, D. Avila, S. Abdalla, C. Willing, and J. Development Team. 2016. Jupyter Notebooks-a publishing format for reproducible computational workflows. Positioning and Power in Academic Publishing: Players, Agents and Agendas. 87–90. doi:10.3233/978-1-61499-649-1-87.

Kortlever, R.M., N.M. Sodir, and C.H. Wilson. 2017. Myc Cooperates with Ras by Programming Inflammation and Immune Suppression In Brief Oncogenic Myc activity orchestrates an immune suppressive tumor microenvironment. Cell. 171. doi:10.1016/j.cell.2017.11.013.

Kruspig, B., T. Monteverde, S. Neidler, A. Hock, E. Kerr, C. Nixon, W. Clark, A. Hedley, S. Laing, S.B. Coffelt, J. Le Quesne, C. Dick, K. Vousden, C.P. Martins, and D.J. Murphy. 2018. The ERBB network facilitates KRAS-driven lung tumorigenesis. Sci Transl Med. 10. doi:10.1126/SCITRANSLMED.AAO2565.

Lloyd, C.M., and B.J. Marsland. 2017. Review Lung Homeostasis: Influence of Age, Microbes, and the Immune System. Immunity. 46:549–561. doi:10.1016/j.immuni.2017.04.005.

Loyher, P.-L., P. Hamon, M. Laviron, A. Meghraoui-Kheddar, E. Goncalves, Z. Deng, S. Torstensson, N. Bercovici, C. Baudesson de Chanville, B. Combadière, F. Geissmann, A. Savina, C. Combadière, and A. Boissonnas. 2018. Macrophages of distinct origins contribute to tumor development in the lung. J. Exp. Med. 215:2536–2553. doi:10.1084/jem.20180534.

Mamedov, M.R., A. Scholzen, R. V. Nair, K. Cumnock, J.A. Kenkel, J.H.M. Oliveira, D.L. Trujillo, N. Saligrama, Y. Zhang, F. Rubelt, D.S. Schneider, Y. hsiu Chien, R.W. Sauerwein, and M.M. Davis. 2018. A Macrophage Colony-Stimulating-Factor-Producing γδ T Cell Subset Prevents Malarial Parasitemic Recurrence. Immunity. 48:350–363.e7. doi:10.1016/J.IMMUNI.2018.01.009.

McCowan, J., F. Fercoq, P.M. Kirkwood, W. T’Jonck, L.M. Hegarty, C.M. Mawer, R. Cunningham, A.S. Mirchandani, A. Hoy, D.C. Humphries, G.R. Jones, C.G. Hansen, N. Hirani, S.J. Jenkins, S. Henri, B. Malissen, S.R. Walmsley, D.H. Dockrell, P.T.K. Saunders, L.M. Carlin, and C.C. Bain. 2021. The transcription factor EGR2 is indispensable for tissue-specific imprinting of alveolar macrophages in health and tissue repair. Sci Immunol. 6:2132. doi:10.1126/SCIIMMUNOL.ABJ2132.

McKenzie, D.R., E.E. Kara, C.R. Bastow, T.S. Tyllis, K.A. Fenix, C.E. Gregor, J.J. Wilson, R. Babb, J.C. Paton, A. Kallies, S.L. Nutt, A. Brüstle, M. Mack, I. Comerford, and S.R. McColl. 2017. IL-17-producing γδ T cells switch migratory patterns between resting and activated states. Nat Commun. 8. doi:10.1038/NCOMMS15632.

Misharin AV, Morales-Nebreda L, Reyfman PA, Cuda CM, Walter JM, McQuattie-Pimentel AC, Chen CI, Anekalla KR, Joshi N, Williams KJN, Abdala-Valencia H, Yacoub TJ, Chi M, Chiu S, Gonzalez-Gonzalez FJ L.A.N.T. Gates K, Homan PJ, Soberanes S, Dominguez S, Morgan VK, Saber R, Shaffer A, Hinchcliff M, Marshall SA, Bharat A, Berdnikovs S, Bhorade SM, Bartom ET, Morimoto RI, Balch WE, Sznajder JI, Chandel NS, Mutlu GM, Jain M, Gottardi CJ, Singer BD, Ridge KM, Bagheri N, Shilatifard A, Budinger GRS, and Perlman H. 2017. Monocyte-derived alveolar macrophages drive lung fibrosis and persist in the lung over the life span. J Exp Med. 214:2387–2404. doi:10.1084/jem.20162152.

Mugarza, E., F. van Maldegem, J. Boumelha, C. Moore, S. Rana, M. Llorian Sopena, P. East, R. Ambler, P. Anastasiou, P. Romero-Clavijo, K. Valand, M. Cole, M. Molina-Arcas, and J. Downward. 2022. Therapeutic KRAS G12C inhibition drives effective interferon-mediated antitumor immunity in immunogenic lung cancers. Sci. Adv. 8:8780.

Murphy, D.J., M.R. Junttila, L. Pouyet, A. Karnezis, K. Shchors, D.A. Bui, L. Brown-Swigart, L. Johnson, and G.I. Evan. 2008. Distinct Thresholds Govern Myc’s Biological Output In Vivo. Cancer Cell. 14. doi:10.1016/j.ccr.2008.10.018.

Papadopoulos, K.P., L. Gluck, L.P. Martin, A.J. Olszanski, A.W. Tolcher, G. Ngarmchamnanrith, E. Rasmussen, B.M. Amore, D. Nagorsen, J.S. Hill, and J. Stephenson. 2017. Cancer Therapy: Clinical First-in-Human Study of AMG 820, a Monoclonal Anti-Colony-Stimulating Factor 1 Receptor Antibody, in Patients with Advanced Solid Tumors. Clin Cancer Res. 23:5703–5710. doi:10.1158/1078-0432.CCR-16-3261.

Park, M.D., I. Reyes-Torres, J. Leberichel, P. Hamon, N.M. Lamarche, S. Hegde, M. Belabed, L. Troncoso, J.A. Grout, A. Magen, E. Humblin, A. Nair, M. Molgora, J. Hou, J.H. Newman, A.M. Farkas, A.M. Leader, T. Dawson, D. D’souza, S. Hamel, A. Rodriguez Sanchez-Paulete, B. Maier, N. Bhardwaj, J.C. Martin, A.O. Kamphorst, E. Kenigsberg, M. Casanova-Acebes, A. Horowitz, B.D. Brown, L. Ferrari De Andrade, M. Colonna, T.U. Marron, and M. Merad. 2023. TREM2 macrophages drive NK cell paucity and dysfunction in lung cancer. Nat Immunol. 24:792–801. doi:10.1038/s41590-023-01475-4.

Patel, A.J., P. Nightingale, B. Naidu, M.T. Drayson, G.W. Middleton, and A. Richter. 2020. Characterising the impact of pneumonia on outcome in non-small cell lung cancer: identifying preventative strategies. J Thorac Dis. 12:2236. doi:10.21037/JTD.2020.04.49.

Pizzolato, G., H. Kaminski, M. Tosolini, D.M. Franchini, F. Pont, F. Martins, C. Valle, D. Labourdette, S. Cadot, A. Quillet-Mary, M. Poupot, C. Laurent, L. Ysebaert, S. Meraviglia, F. Dieli, P. Merville, P. Milpied, J. Déchanet-Merville, and J.J. Fournié. 2019. Single-cell RNA sequencing unveils the shared and the distinct cytotoxic hallmarks of human TCRVδ1 and TCRVδ2 γδ T lymphocytes. Proc Natl Acad Sci U S A. 116:11906–11915. doi:10.1073/PNAS.1818488116/-/DCSUPPLEMENTAL.

Rashidi, S., C. Fernández-Rubio, R. Manzano-Román, R. Mansouri, R. Shafiei, M. Ali-Hassanzadeh, A. Barazesh, M. Karimazar, G. Hatam, and P. Nguewa. 2021. Potential therapeutic targets shared between leishmaniasis and cancer. Parasitology. 148:655–671. doi:10.1017/S0031182021000160.

Razak, A.R., J.M. Cleary, V. Moreno, M. Boyer, E. Calvo Aller, W. Edenfield, J. Tie, R. Donald Harvey, A. Rutten, M.A. Shah, A.J. Olszanski, D. Jäger, N. Lakhani, D.P. Ryan, E. Rasmussen, G. Juan, H. Wong, N. Soman, M.-A. Damiette Smit, D. Nagorsen, and K.P. Papadopoulos. 2020. Safety and efficacy of AMG 820, an anti-colony-stimulating factor 1 receptor antibody, in combination with pembrolizumab in adults with advanced solid tumors. J Immunother Cancer. 8:1006. doi:10.1136/jitc-2020-001006.

Ribot, J.C., N. Lopes, and B. Silva-Santos. 2020. γδ T cells in tissue physiology and surveillance. Nature Reviews Immunology 2020 21:4. 21:221–232. doi:10.1038/s41577-020-00452-4.

Silva-Santos, B., S. Mensurado, and S.B. Coffelt. 2019. γδ T cells: pleiotropic immune effectors with therapeutic potential in cancer. Nat Rev Cancer. 19:392–404. doi:10.1038/s41568-019-0153-5.

Street, K., D. Risso, R.B. Fletcher, D. Das, J. Ngai, N. Yosef, E. Purdom, and S. Dudoit. 2018. Slingshot: Cell lineage and pseudotime inference for single-cell transcriptomics. BMC Genomics. 19:1–16. doi:10.1186/S12864-018-4772-0/FIGURES/5.

Subramanian, A., P. Tamayo, V.K. Mootha, S. Mukherjee, B.L. Ebert, M.A. Gillette, A. Paulovich, S.L. Pomeroy, T.R. Golub, E.S. Lander, and J.P. Mesirov. 2005. Gene set enrichment analysis: A knowledge-based approach for interpreting genome-wide expression profiles. PNAS. 102:15545–15550.

Sunaga, S., K. Maki, Y. Komagata, K. Ikuta, and J.-I. Miyazaki. 1997. Efficient removal ofloxP-flanked DNA sequences in a gene-targeted locus by transient expression of Cre recombinase in fertilized eggs. Mol Reprod Dev. 46:109–113. doi:10.1002/(SICI)1098-2795(199702)46:2<109::AID-MRD1>3.0.CO;2-U.

Tang, Z., B. Kang, C. Li, T. Chen, and Z. Zhang. 2019. GEPIA2: an enhanced web server for large-scale expression profiling and interactive analysis. Nucleic Acids Res. 47:W556–W560. doi:10.1093/nar/gkz430.

Wands, J.M., C.L. Roark, M.K. Aydintug, N. Jin, Y.-S. Hahn, L. Cook, X. Yin, J. Dal Porto, M. Lahn, D.M. Hyde, E.W. Gelfand, R.J. Mason, R.L. O’Brien, and W.K. Born. 2005. Distribution and leukocyte contacts of gammadelta T cells in the lung. J Leukoc Biol. 78:1086–1096. doi:10.1189/JLB.0505244.

Weng, R.R., H.-H. Lu, C.-T. Lin, C.-C. Fan, R.-S. Lin, T.-C. Huang, S.-Y. Lin, Y.-J. Huang, Y.-H. Juan, Y.-C. Wu, Z.-C. Hung, C. Liu, X.-H. Lin, W.-C. Hsieh, T.-Y. Chiu, J.-C. Liao, Y.-L. Chiu, S.-Y. Chen, C.-J. Yu, and H.-C. Tsai. 2021. Epigenetic modulation of immune synaptic-cytoskeletal networks potentiates γδ T cell-mediated cytotoxicity in lung cancer. Nat Commun. 12. doi:10.1038/s41467-021-22433-4.

Wingett, S.W., and S. Andrews. 2018. FastQ Screen: A tool for multi-genome mapping and quality control. F1000Res. 7. doi:10.12688/f1000research.15931.1.

Wu, Y., D. Biswas, I. Usaite, M. Angelova, S. Boeing, T. Karasaki, S. Veeriah, J. Czyzewska-Khan, C. Morton, M. Joseph, S. Hessey, J. Reading, A. Georgiou, M. Al-Bakir, N.J. Birkbak, G. Price, M. Khalil, K. Kerr, S. Richardson, H. Cheyne, T. Cruickshank, G.A. Wilson, R. Rosenthal, H. Aerts, M. Hewish, G. Anand, S. Khan, K. Lau, M. Sheaff, P. Schmid, L. Lim, J. Conibear, R. Schwarz, T.L. Kaufmann, M. Huska, J. Shaw, J. Riley, L. Primrose, D. Fennell, A. Hackshaw, Y. Ngai, A. Sharp, O. Pressey, S. Smith, N. Gower, H. Kaur Dhanda, K. Chan, S. Chakraborty, K. Litchfield, K. Thakkar, J. Tugwood, A. Clipson, C. Dive, D. Rothwell, A. Kerr, E. Kilgour, F. Morgan, M. Kornaszewska, R. Attanoos, H. Davies, K. Baker, M. Carter, C.R. Lindsay, F. Gomes, F. Blackhall, L. Priest, M.G. Krebs, A. Chaturvedi, P. Oliveira, Z. Szallasi, G. Royle, C. Veiga, M. Skrzypski, R. Salgado, M. Diossy, A. Kirk, M. Asif, J. Butler, R. Bilancia, N. Kostoulas, M. Thomas, M. MacKenzie, M. Wilcox, A. Nakas, S. Rathinam, R. Boyles, M. Tufail, A. Bajaj, K. Ang, M. Fiyaz Chowdhry, M. Shackcloth, J. Asante-Siaw, A. Leek, N. Totten, J. Davies Hodgkinson, P. Van Loo, W. Monteiro, H. Marshal, et al. 2022. A local human Vδ1 T cell population is associated with survival in nonsmall-cell lung cancer. Nat Cancer. 3:696–709. doi:10.1038/s43018-022-00376-z.

