## Supplemental figures for "γδ T cells impair airway macrophage differentiation in lung adenocarcinoma"

**Supplemental material**:

**Video 1. Intravascular γδ T cells.** 3D view of lung parenchyma. First zoom-in shows a $CD27^-$ γδ T cell and, the second zoom-in, a $CD27^+$ γδ T cell. Vasculature (CD31, green), collagen (second harmonic generation, glow palette), leukocytes (CD45, cyan), macrophages (CD68, orange), CD27 (red), and TCRδ (magenta).

**Video 2. Infiltrating γδ T cells.** 3D view of lung tumour. Zoom-in featuring a $CD27^+$ γδ T cell within the tumour, adjacent to the vasculature. Vasculature (CD31, green), collagen (second harmonic generation, glow palette), leukocytes (CD45, cyan), macrophages (CD68, orange), CD27 (red), and TCRδ (magenta).

**Video 3. Interacting γδ T cells.** 3D view of lung tumour-associated adventitial cuff. Zoom-in featuring a $CD27^+$ γδ T cell in contact with a macrophage. Vasculature (CD31, green), collagen (second harmonic generation, glow palette), leukocytes (CD45, cyan), macrophages (CD68, orange), CD27 (red), and TCRD (magenta).

**Video 4. Long-lasting γδ T cells interactions with airway macrophages.** Time-lapse on a precision-cut lung slice of tumour margin imaged every 95.5 seconds. Zoom-ins featuring γδ T cells interacting with airway macrophages. Vasculature (CD31, green), airway macrophages (CD11c, orange), TCRδ (magenta), and Hoescht (blue).

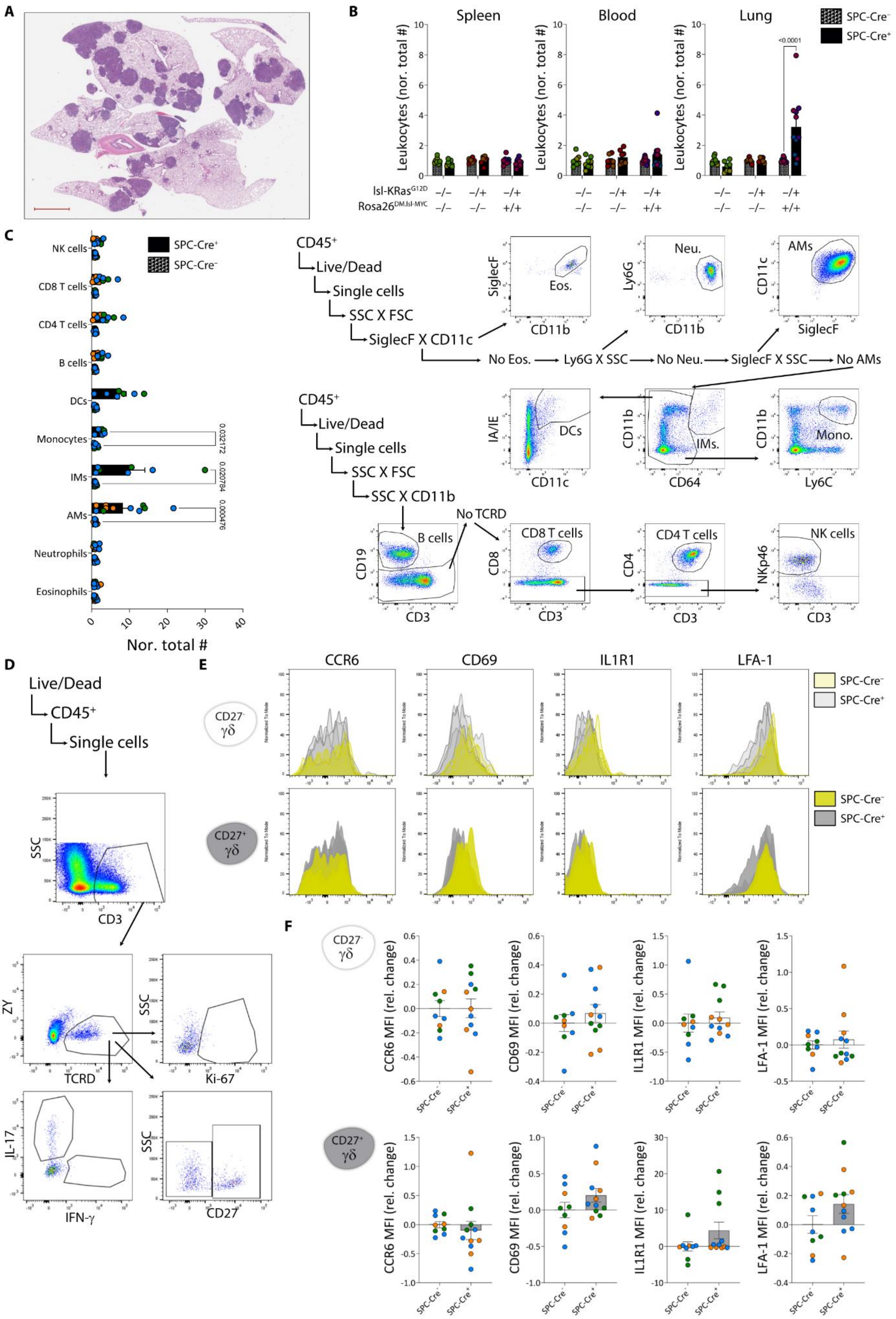
A
B
Spleen
Blood
Lung
SPC-Cre⁻
SPC-Cre⁺
<0.0001
Leukocytes (nor. total #)
lsl-KRas G12D
Rosa26 DM.lsl-MYC
C
NK cells
CD8 T cells
CD4 T cells
B cells
DCs
Monocytes
IMs
AMs
Neutrophils
Eosinophils
Nor. total #
0.032172
0.020784
0.000476
CD45⁺
Live/Dead
Single cells
SSC X FSC
SiglecF X CD11c
No Eos.
Ly6G X SSC
No Neu.
SiglecF X SSC
No AMs
Eos.
Neu.
AMs
SSC X CD11b
No TCRD
DCs
IMs.
Mono.
B cells
CD8 T cells
CD4 T cells
NK cells
D
Live/Dead
CD45⁺
Single cells
E
CCR6
CD69
IL1R1
LFA-1
CD27⁻ γδ
CD27⁺ γδ
F
CCR6 MFI (rel. change)
CD69 MFI (rel. change)
IL1R1 MFI (rel. change)
LFA-1 MFI (rel. change)

**Figure S1. Immune phenotype of KM mice. (A)** Representative H&E of SPC-Cre$^{+}$ KM mouse lungs 8 weeks after allele activation. Scale bar: 2 mm. **(B)** Normalised total number of leukocytes in spleen, blood, and lungs for different genotypes. Data were analysed by two-way ANOVA followed by Sidak's post-test. **(C)** Main leukocyte populations in lungs of KM mice (SPC-Cre$^{-}$ and SPC-Cre$^{+}$). Data were analysed by Mann-Whitney test with Holm-Sidak correction for multiple comparisons. Schematic gating strategy on the right. **(D)** Schematic gating strategy for γδ T cells. **(E)** Histograms of gated pulmonary CD27$^{-}$ and CD27$^{+}$ γδ T cells for CCR6, CD69, IL1R1, and LFA-1, from one representative experiment (related to **Fig. 1C**). **(F)** Gated on blood CD27$^{-}$ (white bars) and CD27$^{+}$ (grey bars) γδ T cells, MFI relative change for CCR6, CD69, IL1R1, and LFA-1. **(B, C, F)** Each dot represents a mouse, coloured by independent experiment. Data are presented as mean ± SEM.

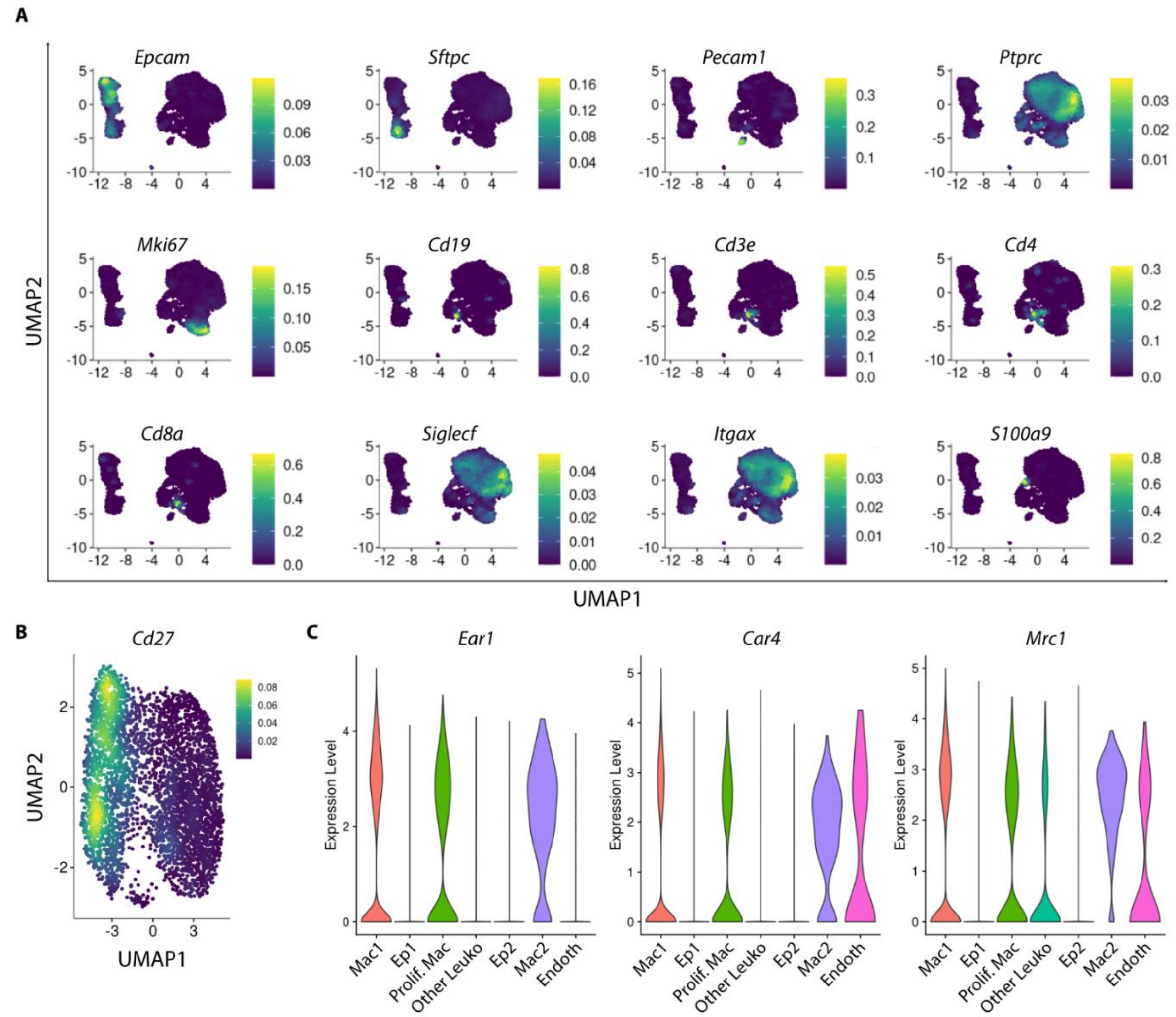


**Figure S2. Genes used to annotate scRNA-seq clusters. (A)** Density plots showing expression of *Epcam*, *Sftpc*, *Pecam1*, *Ptprc*, *Mki67*, *Cd19*, *Cd3e*, *Cd4*, *Cd8a*, *Siglecf*, *Itgax*, and *S100a9* on UMAP dimensionality reduction for scRNA-seq data from microdissected SPC-Cre$^{+}$ KM lung tumours. **(B)** Density plot showing expression of *Cd27* on UMAP dimensionality reduction for scRNA-seq data from naïve pulmonary $\gamma\delta$ T cells (Edwards et al., 2023). **(C)** Violin plots showing the expression of *Ear1*, *Car4*, and *Mrc1* by cluster in the scRNA-seq data from microdissected SPC-Cre$^{+}$ KM lung tumours.

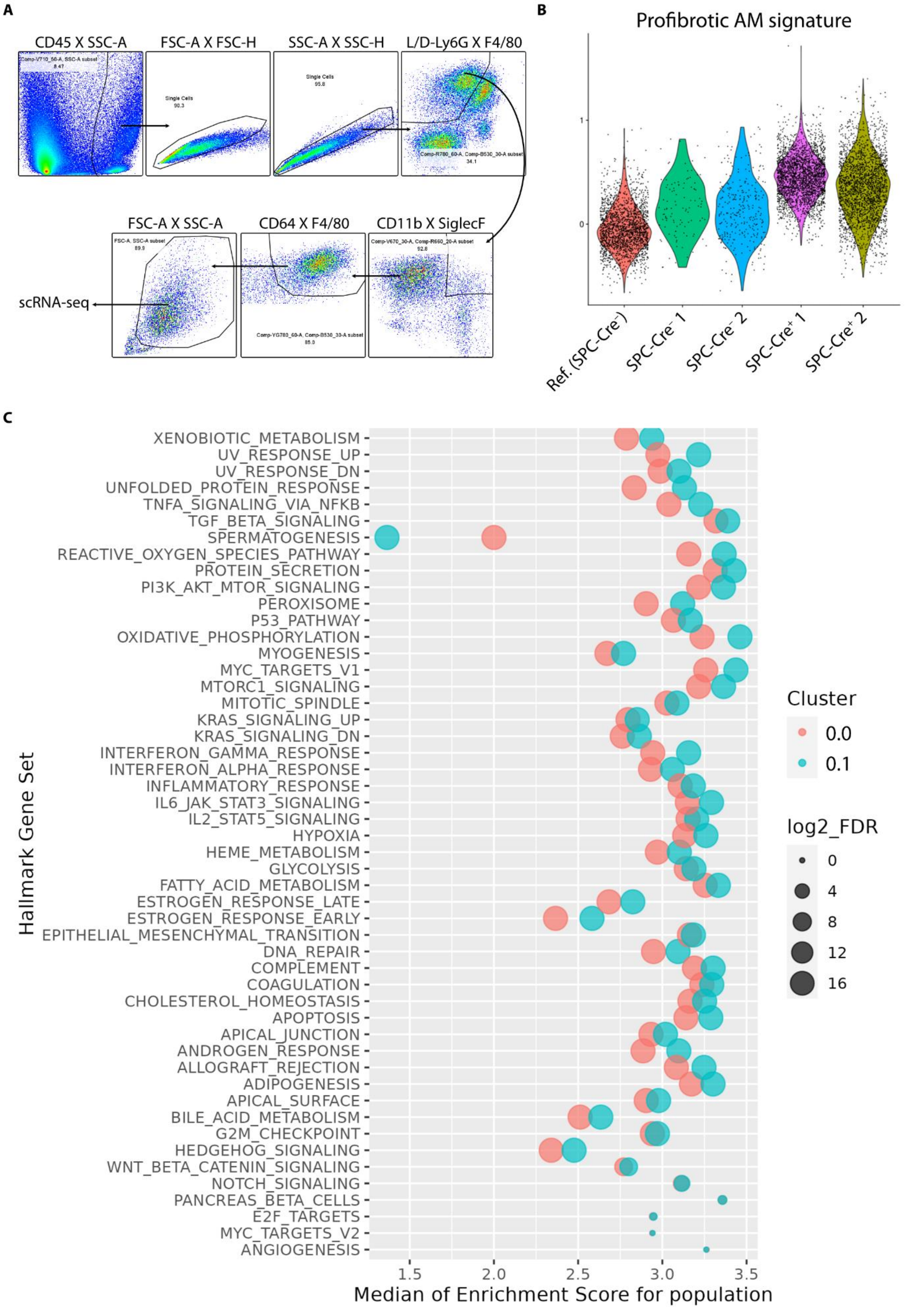
A
CD45 X SSC-A
FSC-A X FSC-H
SSC-A X SSC-H
L/D-Ly6G X F4/80
FSC-A X SSC-A
CD64 X F4/80
CD11b X SiglecF
scRNA-seq
B
Profibrotic AM signature
1
0
Ref. (SPC-Cre-)
SPC-Cre- 1
SPC-Cre- 2
SPC-Cre+ 1
SPC-Cre+ 2
C
Hallmark Gene Set
XENOBIOTIC_METABOLISM
UV_RESPONSE_UP
UV_RESPONSE_DN
UNFOLDED_PROTEIN_RESPONSE
TNFA_SIGNALING_VIA_NFKB
TGF_BETA_SIGNALING
SPERMATOGENESIS
REACTIVE_OXYGEN_SPECIES_PATHWAY
PROTEIN_SECRETION
PI3K_AKT_MTOR_SIGNALING
PEROXISOME
P53_PATHWAY
OXIDATIVE_PHOSPHORYLATION
MYOGENESIS
MYC_TARGETS_V1
MTORC1_SIGNALING
MITOTIC_SPINDLE
KRAS_SIGNALING_UP
KRAS_SIGNALING_DN
INTERFERON_GAMMA_RESPONSE
INTERFERON_ALPHA_RESPONSE
INFLAMMATORY_RESPONSE
IL6_JAK_STAT3_SIGNALING
IL2_STAT5_SIGNALING
HYPOXIA
HEME_METABOLISM
GLYCOLYSIS
FATTY_ACID_METABOLISM
ESTROGEN_RESPONSE_LATE
ESTROGEN_RESPONSE_EARLY
EPITHELIAL_MESENCHYMAL_TRANSITION
DNA_REPAIR
COMPLEMENT
COAGULATION
CHOLESTEROL_HOMEOSTASIS
APOPTOSIS
APICAL_JUNCTION
ANDROGEN_RESPONSE
ALLOGRAFT_REJECTION
ADIPOGENESIS
APICAL_SURFACE
BILE_ACID_METABOLISM
G2M_CHECKPOINT
HEDGEHOG_SIGNALING
WNT_BETA_CATENIN_SIGNALING
NOTCH_SIGNALING
PANCREAS_BETA_CELLS
E2F_TARGETS
MYC_TARGETS_V2
ANGIOGENESIS
1.5
2.0
2.5
3.0
3.5
Median of Enrichment Score for population
Cluster
0.0
0.1
log2_FDR
0
4
8
12
16

**Figure S3. scRNA-seq of macrophages from microdissected SPC-Cre$^{-}$ and SPC-Cre$^{+}$ KM lungs. (A)** Gating strategy used for the isolation of macrophages from microdissected SPC-Cre$^{-}$ and SPC-Cre$^{+}$ KM lungs for scRNA-sequencing. **(B)** Violin plot showing the expression of a consensus profibrotic AM signature (Joshi et al., 2020) by sample. **(C)** Median of Enrichment Score for each of the Gene Set Enrichment Analysis Hallmark gene sets for the two tumour-associated AM clusters (related to **Fig. 3D**).
